# Biofunctional nanodot arrays in living cells uncover synergistic co-condensation of Wnt signalodroplets

**DOI:** 10.1101/2022.06.11.495600

**Authors:** Michael Philippi, Christian P. Richter, Marie Kappen, Isabelle Watrinet, Yi Miao, Mercedes Runge, Lara Jorde, Sergej Korneev, Michael Holtmannspötter, Rainer Kurre, Joost C. M. Holthuis, K. Christopher Garcia, Andreas Plückthun, Martin Steinhart, Jacob Piehler, Changjiang You

## Abstract

Qualitative and quantitative analysis of transient signaling platforms in the plasma membrane has remained a key experimental challenge. Here, we have developed biofunctional nanodot arrays (bNDAs) to spatially control dimerization and clustering of cell surface receptors at nanoscale. High-contrast bNDAs with spot diameters of ∼300 nm were obtained by capillary nanostamping of BSA bioconjugates, which were subsequently biofunctionalized by reaction with tandem anti-GFP clamp fusions. We achieved spatially controlled assembly of active Wnt signalosomes at the nanoscale in the plasma membrane of live cells by capturing the co-receptor Lrp6 into bNDAs via an extracellular GFP tag. Strikingly, we observed co-recruitment of co-receptor Frizzled-8 as well as the cytosolic scaffold proteins Axin-1 and Disheveled-2 into Lrp6 nanodots in the absence of ligand. Density variation and the high dynamics of effector proteins uncover highly cooperative liquid-liquid phase separation (LLPS)-driven assembly of Wnt “signalodroplets” at the plasma membrane, pinpointing the synergistic effects of LLPS for Wnt signaling amplification. These insights highlight the potential of bNDAs for systematically interrogating nanoscale signaling platforms and condensation at the plasma membrane of live cells.

## Introduction

The plasma membrane of mammalian cells is dynamically compartmentalized at the micro- and nanoscale, caused by a hierarchical interplay of corral formation by the cortical actin meshwork and segregation by protein-lipid interactions in conjunction with phase transitions of proteins and lipids (1–5). The resulting hierarchical compartmentalization in the plasma membrane intricately regulates the functional organization of signaling complexes (6–9), in particular by promoting dynamic receptor clustering, which has emerged as a key principle for amplifying downstream signaling (10–14). The molecular and cellular mechanisms responsible for receptor clustering, however, have remained largely unresolved (11). Recently, liquid-liquid phase separation (LLPS) of proteins induced by multivalent protein-protein interactions emerged as a key principle driving dynamic receptor clustering (15–21). Experimentally pinpointing and characterizing the involvement of LLPS in the formation and activation of signaling complexes in the plasma membrane has remained challenging due to the nanoscale dimensions and the high dynamics of such entities.

Here, we introduce biofunctional nanodot array (bNDA) technology as a generic approach to assemble signaling platforms with well-defined geometry in the plasma membrane of live cells. To this end, we employed capillary nanostamping (22, 23) for printing functionalized bovine serum albumin (BSA) onto hydrophobic glass coverslips, yielding nanodots with < 300 nm diameter. By subsequent conjugation with an engineered GFP binder, efficient, high-density capturing of GFP-tagged proteins was achieved, yielding very high contrast in *vitro* and in live cells. For proof-of-concept experiments, we employed bNDAs to assemble functional Wnt signalosomes (24) that have recently been proposed to involve LLPS (25, 26). The canonical Wnt signaling pathway activates the transcription factor β-catenin, which is an evolutionally highly conserved regulator of stem cell fate decision (27–29). Wnt signalosome formation is initiated by crosslinking of low-density lipoprotein receptor-related protein 5/6 (Lrp5/6) and the co-receptor Frizzled (Fzd) in the plasma membrane via the Wnt ligand (30) (Fig. 1A). Receptor heterodimerization initiates recruitment of the cytosolic scaffold proteins Axin and Disheveled (Dvl), leading to the formation of multiprotein assemblies in the plasma membrane, the so-called Wnt signalosomes (24, 31).

**Fig. 1.**
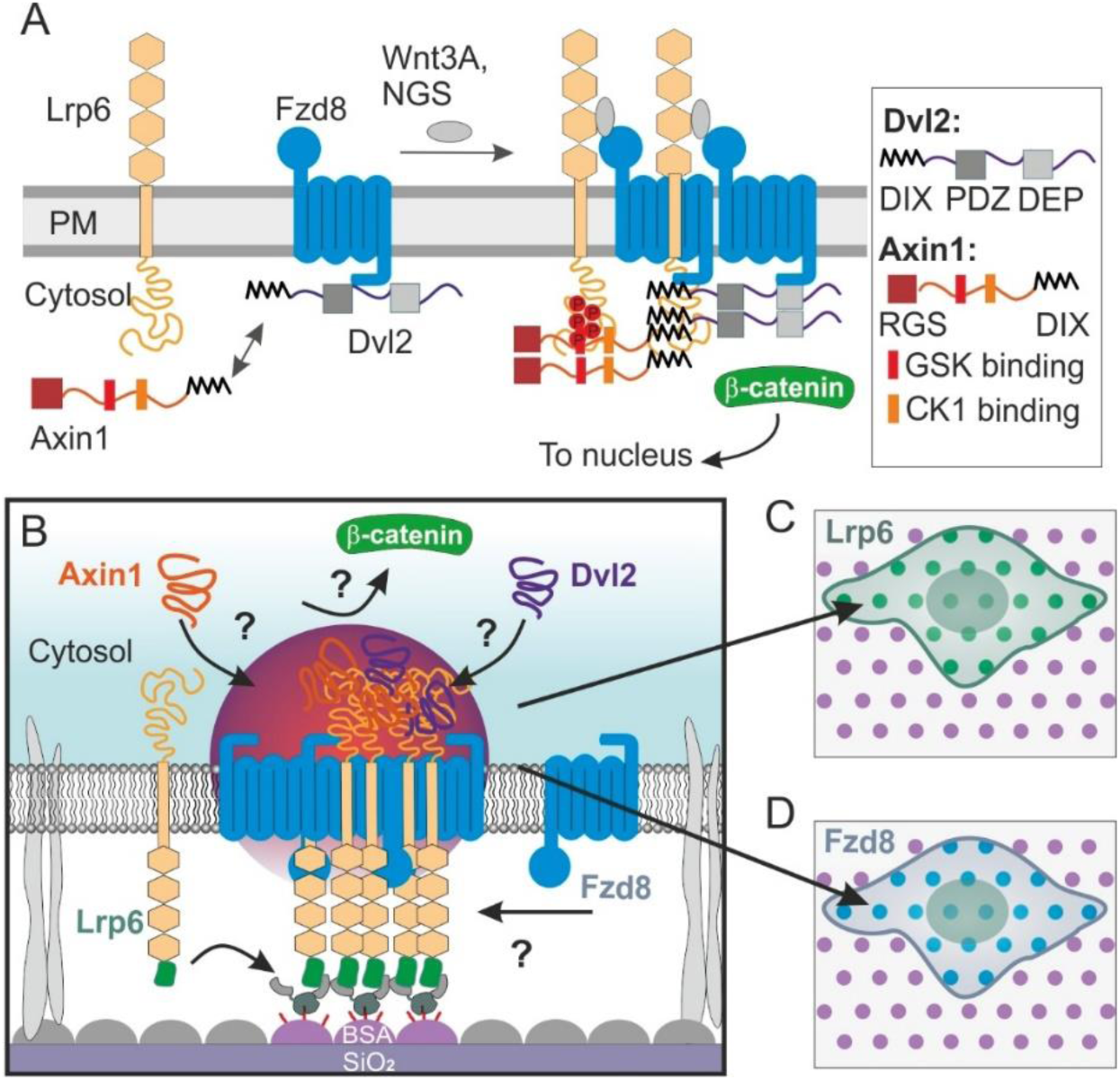
Reorganization of the Wnt signaling relevant proteins in live cell plasma membrane by nanodot arrays (NDAs). (A) Scheme summarizing the current view Wnt signalosome formation responsible for activating the Wnt/β-catenin pathway. The intrinsically disordered scaffold proteins Dvl2 and Axin1 and their binding motifs are outlined in the box. (B) Concept of live cell protein nanodot arrays for inducing Wnt signalosome assembly by locally clustering Lrp6 in the plasma membrane via an extracellular GFP-tag. (C, D) Potential recruitment of Wnt signaling proteins labeled Fzd8, Axin1 and Dvl2 into Lrp6 bNDAs can be detected and quantified by TIRF microscopy.

The molecular mechanisms underlying Wnt signalosome formation, however, are currently under debate. Interestingly, not only ligand-induced heterodimerization and/or hetero-oligomerization of Fzd and Lrp6 can initiate signalosome formation and β-catenin activation (32–37), but also homo-oligomerization of Lrp6 (38, 39), with its intracellular multiple binding motifs that interact with Axin1 playing a critical role (40, 41). These observations could be explained by LLPS being a driving force to form Wnt signalsome as condensates (25, 26), and indeed, co-condensates of the cytosolic scaffold proteins Axin and Disheveled (Dvl) have been observed (42–44). However, systematic analyses of Wnt signalosome assembly and dynamics in the plasma membrane under well-defined conditions have been lacking so far.

Here, we applied bNDAs for spatially controlling Wnt signalosome assembly in plasma membrane of live cells (Fig. 1B). Strikingly, dimerizing and clustering of Lrp6 in bNDAs promoted ligand-independent recruitment of the co-receptor Fzd8 and the cytosolic effector proteins Axin1 and Dvl2 as well as activation of downstream signaling. Quantitative analysis at single nanodot level revealed dynamic, highly cooperative effector binding and a critical role of Lrp6 density. Our insights strongly support that Wnt signalsome assembly is driven by co-condensation of Axin and Dvl, which is regulated by local enrichment of Axin upon transient interaction with clustered Lrp6.

## Results

### Protein-based surface functionalization for contact lithography with nanoscale resolution

To achieve efficient, high-density surface biofunctionalization and high precision capillary nanostamping, we applied bovine serum albumin (BSA) as an ink, which has proven well compatible with high-contrast contact lithography via physisorption onto hydrophilic and hydrophobic surfaces (45, 46). For subsequent functionalization, BSA was conjugated with amine-functionalized HaloTag ligand (HTL) (47) using amide coupling chemistry. The synthesized ^HTL^BSA had an average degree of functionalization of 13 HTL moieties per BSA according to mass spectrometry and was used for bioorthogonal covalent capturing of target proteins fused to the HaloTag (47) as depicted in Fig. 2A. To promote adsorption rates of BSA to glass surfaces and to minimize spreading of the printed solution, the glass substrates were rendered hydrophobic by silanization. Surface modification with 1-napthylmethyltrichlorosilane (NMTS) turned out most efficient compared to other hydrophobic silanes (Supplementary Fig. S1A), for which the binding kinetics of BSA was quantified by label-free detection based on reflectance interference (RIF) under flow-through conditions (48–50). Very rapid and stable BSA coating was observed on NMTS-treated silica substrates (Fig. 2B, Fig. S1b) with the final mass signal of ∼3.5 ng/mm^2^ being in good agreement with formation of a BSA monolayer (51). Specific immobilization of the mEGFP-HaloTag fusion protein on the ^HTL^BSA-coated surface was confirmed by real-time total internal reflection fluorescence (TIRF) detection (Fig. 2C), with an association rate constant (*k_on_*) of ∼10^3^ M^-1^s^-1^ (Supplementary Table S1) in line with previous kinetics studies of the HaloTag-HTL interaction (52).

**Fig. 2.**
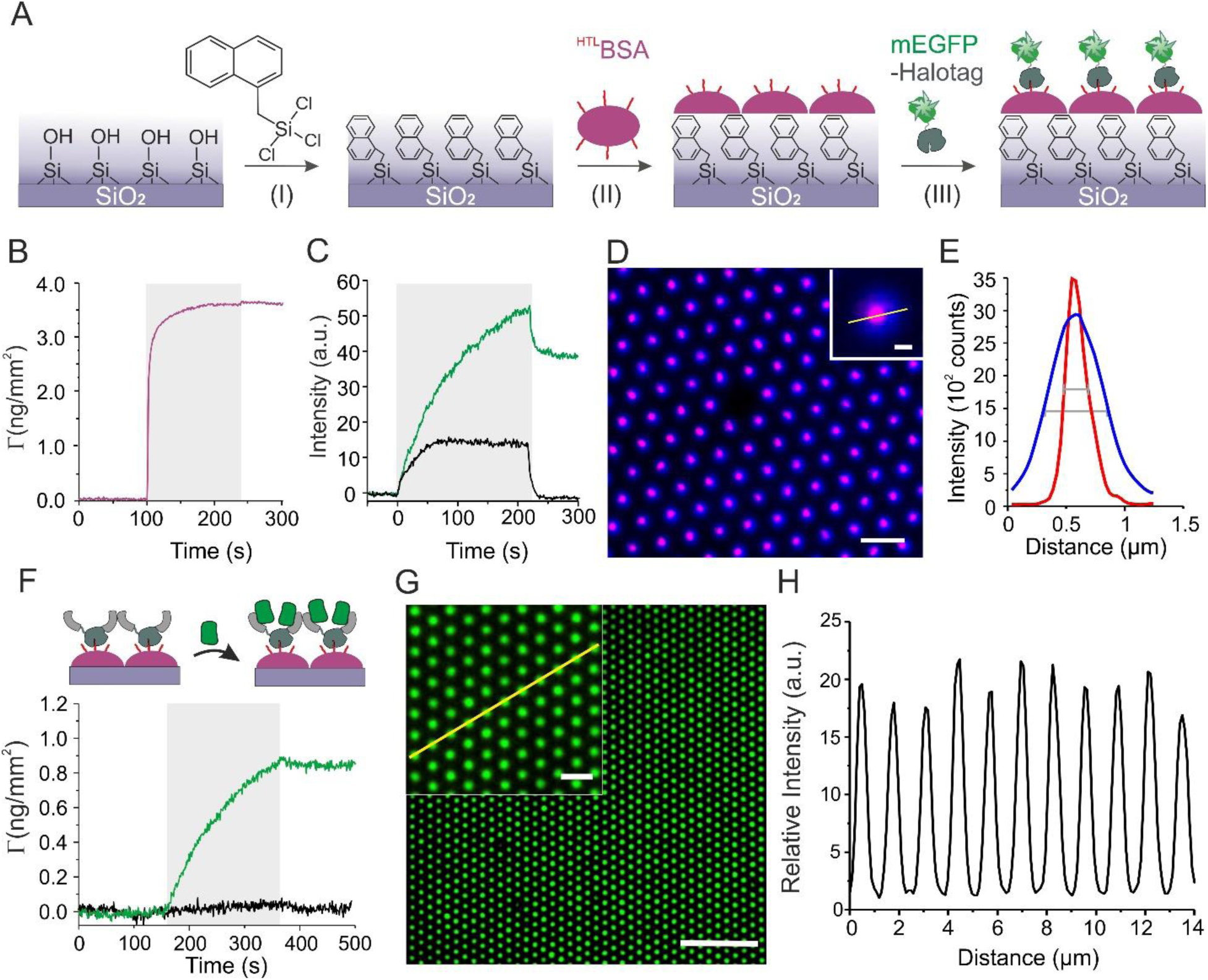
Surface functionalization for fabricating high-contrast bNDA on glass support. (A) Scheme of the procedure: silanization by 1-napthylmethyltrichlorosilane (NMTS) for rendering the surface hydrophobic (I), binding of HaloTag Ligand-BSA conjugate (^HTL^BSA) to NMTS-coated substrates (II), and specific biofunctionalization with HaloTag-mEGFP. (B) Real-time kinetics of 10 µM ^HTL^BSA binding onto a NMTS-coated silica surface monitored by label-free RIF detection. (C) Fluorescence signal during injection of 2 µM Halo-mEGFP binding to a ^HTL^BSA-coated substrate (green) as detected by TIRFS. Only the background signal was detected upon injection of 2 µM mEGFP without HaloTag on the same surface (black). (D) Merged dSTORM image (red) and diffraction limited, unprocessed TIRF image (blue) of ^AT647N^BSA printed into an bNDA. Scale bar: 1 µm (inset: 100 nm). (E) Intensity profiles across the line shown in panel (D). The diameters of nanodots were obtained as the FWHM of the intensity profile. (F) mEGFP binding after functionalization of ^HTL^BSA-coated surface with tdClamp-HaloTag as detected by RIF. (G) Deconvolved TIRF microscopy image of mEGFP bound to tdClamp-functionalized bNDA. Scale bar: 10 µm, inset: 2 µm. (H) Relative intensity profile highlighted by the yellow line shown in the inset of panel G.

To explore the capability of generating high-contrast biofunctional nanodot arrays (bNDAs), BSA labeled with ATTO647N (^AT647N^BSA) was stamped onto NMTS-treated glass microscopy coverslips using an established protocol (22). Because the dot sizes and distance are close to the diffraction limit (Fig. S1C), we applied image deconvolution for more reliable image analysis (22), which not only increased the resolution, but also the contrast (Fig. S1D). To more precisely quantify the diameter of the nanodots, we resolved the spatial distribution of printed BSA in nanodots beyond the diffraction limit by direct stochastic optical reconstruction microscopy (dSTORM) of the printed ^AT647N^BSA bNDAs (Fig. 2D, E, Fig. S1E). The dSTORM super-resolution images revealed an average diameter of 256 ± 33 nm (*N* = 10) for the nanodots, which is substantially below the 450 ± 68 nm (*N* = 11) observed for deconvolved TIRFM images. The contrast of bNDA was calculated as the central intensity of nanodot divided by the intensity baseline between nanodots. In line with the increased resolution, the dSTORM images moreover revealed the highest contrast achieved by BSA capillary nanoprinting as the consequence of background-free imaging between the nanodots (Fig. S1H).

### Efficient capturing of GFP-tagged proteins into bNDAs

Staining of ^HTL^BSA bNDAs with HaloTag-mEGFP yielded a good contrast of 10 ± 2.6 (N=15), but the intensities of dots within nanopatterns showed a rather heterogeneous distribution (Fig. S2A-C). Similarly, direct capturing of GFP-tagged receptors, also carrying a HaloTag, in the plasma membrane of live cells lacked efficiency and homogeneity, yielding a contrast below 10 (Fig. S2D-F). Ascribing this performance to the relatively slow reaction of the HaloTag with ^HTL^BSA (cf. Fig. 2C and Table S1), we turned to a robust, high-affinity GFP binder, the so-called “GFP-clamp” engineered from DARPins (53) for capturing GFP-tagged proteins into bNDAs. To furthermore promote dimerization of captured receptors at molecular scale, which has been implicated in the activation of Wnt receptors (35, 36), we fused two copies of the GFP-clamp via a linker that included a HaloTag (tdClamp-HaloTag) for in *situ* coupling to immobilized ^HTL^BSA. Rapid and specific binding of mEGFP to ^HTL^BSA-coated surfaces functionalized with tdClamp-HaloTag was confirmed by reflectance interference detection (Fig. 2F) yielding a 25-fold higher *k_on_* than found for direct binding of HaloTag to ^HTL^BSA (Supplementary Table S1). From the signal amplitude, 1/3 of a protein monolayer was estimated to have formed, which corresponds to a binding capacity of ∼13,000 GFP molecules/µm². As a result of this improved capturing strategy, binding of 200 nM mEGFP to ^HTL^BSA bNDAs pretreated with tdClamp-HaloTag robustly yielded a strikingly high contrast of 19.4 ± 1.9 (N = 9) (Fig. 2G,H), providing ideal conditions for receptor clustering in live cells.

### Efficient reorganization of Lrp6 in the plasma membrane induces active Wnt signalosomes

We employed tdClamp-functionalized bNDAs for exploring whether Wnt signalosome assembly could be triggered by clustering Lrp6 as depicted in Fig. 3A. To this end, HeLa cells co-expressing Lrp6 fused to an N-terminal mEGFP-tag and Frizzled 8 fused to the SNAP-tag (SNAP-Fzd8) labeled with Dy647 were cultured on tdClamp-functionalized bNDAs. Strong enrichment of mEGFP-Lrp6 within nanodots was observed, corroborating that highly efficient and selective immobilization of transmembrane receptor was induced (Fig. 3B). Strikingly, significant co-clustering of Fzd8 in bNDAs was observed (Fig. 3B and Fig. S2J-L), which was strongly dependent on the Lrp6 density (Fig. 3C-E). These experiments highlighted that Fzd8 can interact with clustered Lrp6 in the absence of a heterodimerizing agonist such as Wnt in an Lrp6 density-dependent manner. In the contrast, clustering of mEGFP-fused Fzd8 in bNDAs did not recruit Lrp6 (Fig. S2M-O), indicating that Lrp6 bNDAs might be Wnt signaling relevant instead of the Fzd8 bNDAs.

**Fig. 3.**
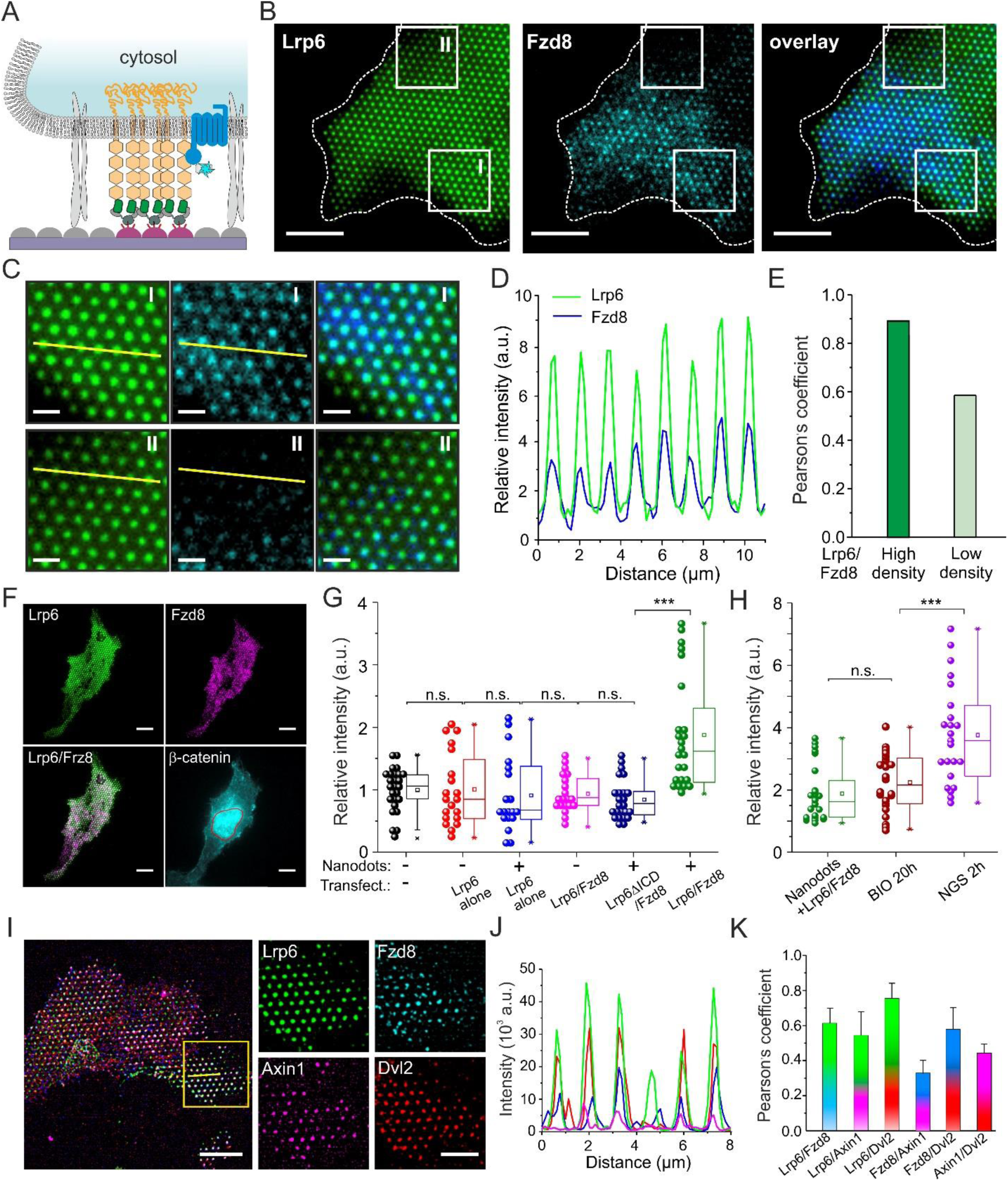
Assembly of functional Wnt signalosomes upon clustering Lrp6 into bNDAs. (A) Scheme illustrating the enrichment of Lrp6 into bNDA via an N-terminal mEGFP-tag. Signalosome assembly was probed by co-expression of Fzd8 fused to an N-terminal SNAP-tag labeled with Dy647. (B) Representative TIRF microscopy image of a live HeLa cell co-expressing mEGFP- Lrp6 (green) and ^DY647^SNAP-Fzd8 (cyan) cultured on bNDA. Scale bars: 10 µm. (C) Zoom in regions with higher (I) and lower (II) density of captured Lrp6. Scale bars: 2 µm. (D) Relative intensity profile marked in panel C region I. (E) Pearson’s correlation coefficient of the marked regions I and II. (F) Representative TIRF microscopy images of a fixed HeLa cell expressing mEGFP-Lrp6 (green) and SNAP-Fzd8 labeled with Dy549 (magenta) captured into an bNDA. After fixation, β-catenin was stained by immunofluorescence and imaged by epifluorescence microscopy (cyan). Scale bars: 10 µm. (G, H) β-catenin levels in the nucleus of HeLa cells cultured under different conditions quantified by IF in individual cells. Statistics by t-test. p value > 0.05 for non-significant (n.s.). p <0.001 for ***. (I-K) Four-color TIRF microscopy imaging of cytosolic effector proteins Axin1 and Dvl1 in Wnt signalodots. (I) Representative TIRF image of HeLa cell co-expressing mEGFP-Lrp6 (green), SNAP-Fzd8 (magenta), mTagBFP-Axin1 (blue) and Dvl2- td-mCherry (red). Overview images with all four colors overlaid (left) and a zoomed region show all four channels. Scale bars: 10 µm (left), 5 µm (zoomed images). (J) Intensity profile of highlighted line profile in panel (I). (K) Co-localization analysis for all channels.

Since oligomerization of Lrp6 has previously been shown to activate Wnt/β-catenin signaling (38, 39), we speculated that the high-density Lrp6 bNDAs are capable to assemble active signalosomes. To verify Wnt signaling at the single cell level, we quantified the β-catenin density in the nucleus by immunofluorescence (IF) staining (Fig. 3F). For HeLa cells expressing mEGFP- Lrp6 alone cultured on bNDAs, no difference of β-catenin as compared to the basal level in unpatterned, untransfected HeLa was observed (Fig. 3G). However, HeLa cells co-expressing mEGFP-Lrp6 and SNAP-Fzd8 showed ∼2-fold increased β-catenin level in the nucleus upon Lrp6 nanoclustering in bNDAs (Fig. 3G), in good agreement with agonist-activated Wnt signaling (32). By contrast, bNDAs of Lrp6 lacking its intracellular domain (Lrp6ΔICD) did not increase the β- catenin level even in the presence of co-expressed Fzd8 (Fig. 3G), in line with the important tandem PPP(S/T)P motifs located in the Lrp6 ICD being critical for Wnt/β-catenin signaling (38, 54). As a positive control, unpatterned HeLa cells were treated overnight with 1 µM of the glycogen synthase kinase 3 (GSK3) inhibitor BIO (6-bromoindirubin-3-oxime) to prevent phosphorylation of β-catenin and the subsequent degradation. IF resulted to a 2.3±0.9 fold (N=24) increase in the nucleus compared to the basal level (Fig. 3H). A further increase of 3.8±1.6 fold (N=23) was observed for HeLa cells stimulated with the potent next generation surrogate (NGS) Wnt (34) (Fig. 3H). Together, these results confirm that co-clustering of Lrp6 and Fzd8 in nanodot arrays lead to spatially controlled Wnt signalosomes in live cells. Wnt signalosome downstream signaling is mediated by the recruitment of the cytosolic effector proteins Axin and Dvl (55).

Having observed activation of canonical Wnt signaling upon Lrp6 clustering into bNDAs, we therefore also probed co-clustering of Axin1 and Dvl2 in bNDAs. Using four-color TIRF microscopy, strong enrichment of Axin1 and Dvl2 at Lrp6/Fzd8 signalodots could be observed (Fig. 3I-K), corroborating successful assembly of comprehensive and fully functional Wnt signalosomes into bNDAs (termed “Wnt signalodots” in the following).

### Wnt signalosome formation is driven by cytosolic effector proteins

Exploiting the highly controlled assembly of Wnt signalodots, we interrogated the mechanistic principles driving Wnt signalosome formation in more detail. In particular, we were interested in quantifying the interaction and dynamics of Axin1 and Dvl2, respectively. Culturing HeLa cells co-expressing mEGFP-Lrp6, SNAP-Fzd8 and a tandem mCherry-Axin1 fusion (tdmCherry-Axin1) on tdClamp-functionalized bNDAs yielded high co-localization of all three proteins (Fig. 4A-D, Fig. S3A). This observation is in line with previous reports that Axin1 – the core component of the β- catenin destruction complex – is recruited to the plasma membrane during canonical Wnt signaling (54–56). Efficient recruitment of Axin1 to Lrp6 nanodots could be explained by direct interactions of Axin1 to Lrp6 via the tandem PPP(S/T)P motifs of Lrp6 located in its ICD (54, 55). In line with this hypothesis, bNDAs formed from Lrp6 lacking the intracellular domain (Lrp6ΔICD) did not recruit Axin1 (Fig. 4E-H, Fig. S3B). Interestingly, Fzd8 was not enriched by Lrp6ΔICD bNDAs either, suggesting that the intracellular domain of Lrp6 and its interaction with Axin1 plays a key role in Wnt-independent receptor interactions. Similarly, Dvl2 was strongly enriched in Lrp6 bNDAs for experiments carried out in wt HeLa cells that express endogenous Axin1/2 (Fig. 4I-L, Fig. S3C). By contrast, much lower levels of Dvl2 were found for the same experiment carried out in genome engineered HeLa cell line that lacks all Axin variants (HeLa Axin1/2 DKO) (Fig. 4M-P). Strikingly, recruitment of Fzd8 into Lrp6-bNDAs was also substantially diminished under these conditions, corroborating that Axin is involved in mediating co-clustering of the receptor subunits. Moreover, reduced density of Lrp6 was found in these experiments, highlighting the cooperation of receptor and effector interactions in signalosome assembly.

**Fig. 4.**
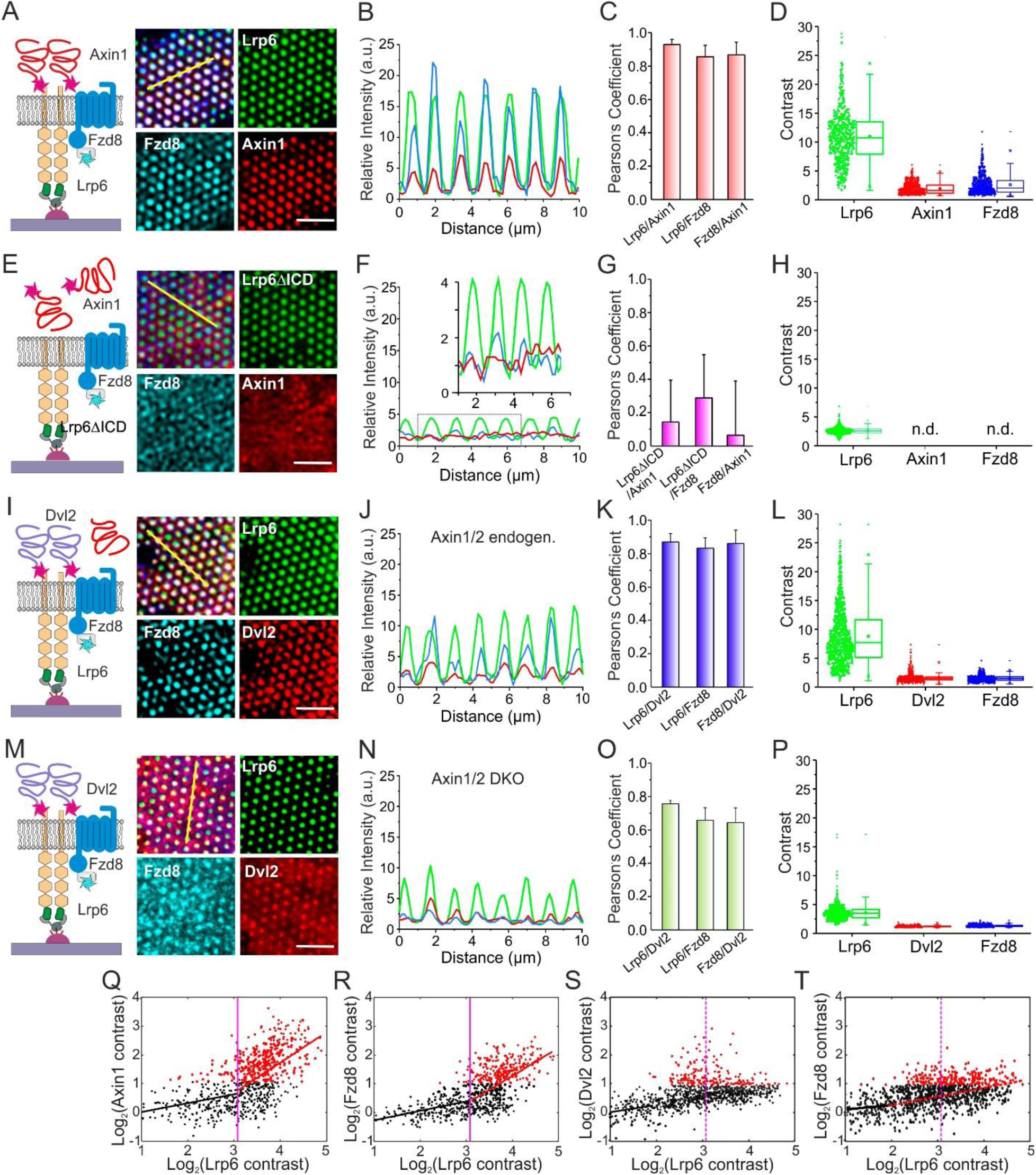
Co-operative recruitment of receptors and effectors into Wnt signalodots. (A-D) Co- recruitment of Axin1 and Fzd8 into Lrp6 bNDAs. (A) Deconvolved 3-color TIRF microscopy images of a HeLa cell co-expressing mEGFP-Lrp6 (green), SNAP-Fzd8 labeled with Dy647 (blue) and tdmCherry-Axin1 (red). (B) Intensity profile highlighted by the yellow line in panel A. (C) Pearson’s correlation analysis with mean±s.d. obtained from 9 line profiles in 3 cells. (D) Single nanodot intensity analysis for all three channels. (E-H) Control experiment with Lrp6 lacking its intracellular domain. (E) Deconvolved TIRF microscopy images of a HeLa cell co-expressing mEGFP-Lrp6ΔICD (green), SNAP-Fzd8 labeled with ^Dy647^ (blue) and tdmCherry-Axin1 (red). (F) Intensity profile highlighted by the yellow line in panel E. (G) Pearson’s correlation analysis with mean ± s.d. obtained from 7 cells. (H) Single nanodot intensity analysis for all three channels. (I- L) Co-recruitment of Dvl2 and Fzd8 into Lrp6 bNDAs. (I) Deconvolved 3-color TIRF microscopy images of a HeLa cell co-expressing mEGFP-Lrp6 (green), SNAP-Fzd8 labeled with ^Dy647^ (blue) and Dvl2-tdmCherry (red). (J) Intensity profile highlighted by the yellow line in panel I. (K) Pearson’s correlation analysis with mean ± s.d. obtained from 9 line profiles in 3 cells. (L) Single nanodot intensity analysis for all three channels. (M-P) Same experiment as shown in panels I-L, but carried out in HeLa cells lacking Axin1 and Axin2 (Axin1/2 DKO). Scale bars: 5 µm for all images. (Q, R) Single nanodot intensity correlation analysis for Lrp6 and Axin1 (Q) and for Lrp6 and Fzd8 (R) from the experiment shown in panel A. (S, T) Single nanodot intensity correlation analysis for Lrp6 and Dvl2 (S) and for Lrp6 and Fzd8 (T) from the experiment shown in panel I.

To pinpoint this feature, we correlated fluorescence intensities of Lrp6 with Axin1 and Fzd8, respectively, in each signalodot. To account for inhomogeneous illumination, the contrasts in each channel were calculated by comparing the intensity of individual dot to its local background. A logarithmic correlation based on ∼700 signalodots confirmed that Axin1 recruitment depends on Lrp6 density in a biphasic mode (Fig. 4Q): At an Lrp6 contrast below 10, Axin1 recruitment remained negligible (contrast < 2). At Lrp6 density above this threshold, however, recruitment of Axin1 strongly increased, reaching a contrast above 10. Recruitment of Fzd8 showed a very similar biphasic dependence on Lrp6 density though with somewhat lower slope of the correlation above the threshold of 10 (Fig. 4R). These biphasic dependence of Lrp6 density supports high cooperativity of receptor and effector interactions, in line with the hypothesis that signalosome formation is being triggered by co-condensation of receptor and effector proteins. Interestingly, biphasic dependence could be observed for neither Dvl2 nor Fzd8 without overexpressing Axin1 (Fig. 4S, T), highlighting the critical role of Axin1 in this process.

### Effector protein dynamics supports LLPS-driven signalodot assembly

Speculating that liquid-liquid phase separation (LLPS) is an important driving force for signalosome formation, we explored the dynamics of Axin1 in signalodot arrays. One hallmark of protein condensates is a rapid exchange with the environment due to transient multivalent interactions (57). We therefore quantified by fluorescence recovery after photobleaching (FRAP) the exchange kinetics at the level of individual signalodots in HeLa cells expressing mEGFP-Lrp6, SNAP-Fzd8 labeled with Dy647 and tdmCherry-Axin1. Strikingly, rapid recovery was observed for Axin1, but not Lrp6 and Fzd8 (Fig. 5A,B, Fig. S4A and Video S1). An average calculated from 30 individual signalodots yielded an overall 34% recovery within 200 s observation time. Likewise, FRAP experiments with HeLa cells expressing mEGFP-Lrp6, SNAP-Fzd8 labeled with Dy647 and Dvl2-tdmCherry revealed dynamic exchange of Dvl2 at the single signalodot level (Fig. 5C,D, Fig. S4B and Video S2).

**Fig. 5.**
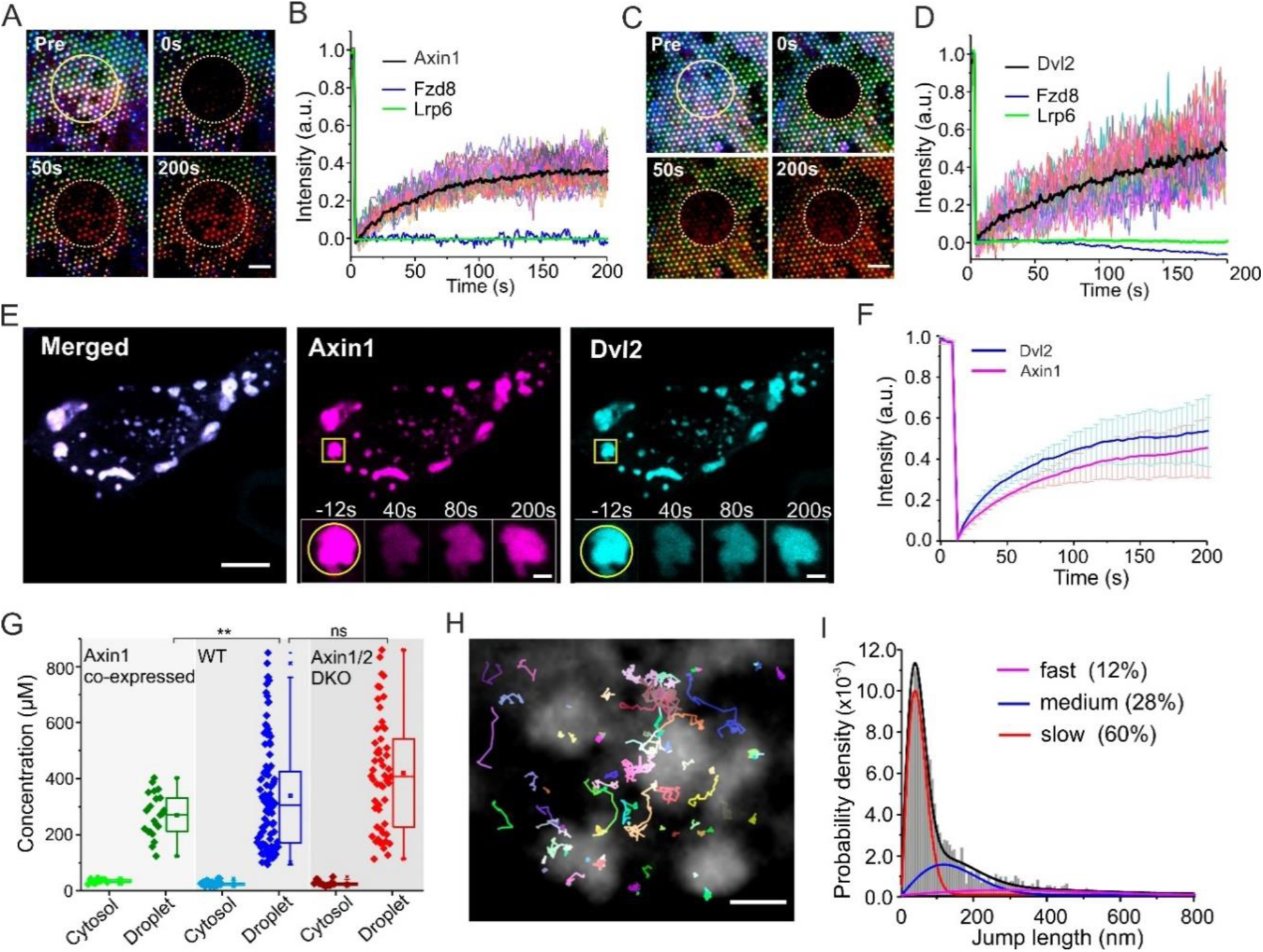
Dynamics of Axin1 and Dvl2 in signalodots and in the cytosol. (A, B) Dynamics of Axin1, Fzd8 and Lrp6 co-recruited into Wnt signalodots. (A) Representative 3-color time-lapse TIRF images of a HeLa cell co-expressing mEGFP-Lrp6 (green), SNAP-Fzd8 labeled with Dy647 (blue) and tdmCherry-Axin1 (red) after photobleaching within the indicated circular region (cf. Video S1). Scale bar: 5 µm. (B) Average FRAP curves of Lrp6 (green), Fzd8 (blue) and Axin1 (black) from the Wnt signalodots. 30 individual recovery curves of Axin1 bNDAs are shown at background. (C, D) Dynamics of tdmCherry-Dvl2, SNAP-Fzd8 and Lrp6 co-recruited into Wnt signalodots. (C) Representative 3-color time-lapse TIRF images of a HeLa cell co-expressing mEGFP-Lrp6 (green), SNAP-Fzd8 labeled with Dy647 (blue) and Dvl2-tdmCherry (red) after photobleaching within the indicated circular region (cf. Video S2). Scale bar: 5 µm. (D) Average FRAP curves of Lrp6 (green), Fzd8 (blue) and Dvl2 (black) from the Wnt signalodots. 30 individual recovery curves of Dvl2 bNDAs are shown. (E, F) Dynamics of Axin1/Dvl2 co-condensates in the cytosol of living cells. (E) Representative confocal microscopy image of HeLa cell overexpressing HaloTag-Axin1 labeled with SiR (magenta) and Dvl2-mCherry (cyan). Inset shows a time-lapse FRAP experiment of a zoomed region (cf. Video S3). Inset scale bar: 2 µm. (F) FRAP curves of Axin1 (magenta) and Dvl2 (blue) obtained from fully bleached droplet (s.d. from triplicate experiments). (G) Quantification of the concentrations of Dvl2-mCherry in the cytosol and in the droplet for three types of HeLa cells: WT, wild type. Axin1/2 DKO: Axin1/2 double knockout HeLa cell. (H, I) Diffusion of Dvl2 inside Dvl2 droplets resolved by single molecule tracking. (H) Colored trajectories of individual Dvl2-HaloTag labeled with SiR in Dvl2-mCherry droplets shown in the background (gray). Scale bar: 2 µm. (I) Step-length histogram analysis of diffusion constants from single molecule trajectories.

As the rapid exchange dynamics is in line with LLPS of Axin1 and Dvl2 in Wnt signalodots, we performed similar FRAP experiments with Axin1/Dvl2 co-condensates that spontaneously formed upon overexpression HaloTag-Axin1 labeled with silicon rhodamine (SiR) and Dvl2-mCherry, respectively, in the cytosol of HeLa cells (Fig. 5E, Video S3). In line with previous observations (42, 44), co-condensates of Axin1 and Dvl2 showed rapid exchange dynamics yielding similar recovery characteristics as in signalodot arrays (Fig. 5F). Dvl2 droplet formation was largely independent of Axin1 expression, but the concentration of Dvl2 in condensates negatively correlated with the Axin1 expression levels (Fig. 5G). The fluidity of droplets was moreover confirmed by sub-droplet FRAP and by observing fusion and fission (Fig. S4D, E and Video S4, Video S5). Single molecule tracking analysis of sub-stoichiometrically labeled Dvl2-HaloTag revealed high mobility inside the droplets with a heterogeneity of the diffusion properties, suggesting different types of interactions (Fig. 5H, I, Video S6). Similar condensation properties were observed for purified Dvl2 recombinantly produced in *E. coli* (Fig. S4F-H).

### Axin1/Dvl2 nanodroplets at the membrane do not trigger signalosome assembly

We therefore hypothesized that Wnt signalosome formation at the plasma membrane is driven by co-condensation of Axin1 and Dvl2 interacting with the co-receptors as depicted in Fig. 6A (38, 55, 58). To explore this hypothesis, we implemented direct recruitment of Axin1 and/or Dvl2 to the plasma membrane using an intracellular anti-GFP nanobody (GFPnb) fused to an artificial transmembrane helix as depicted in Fig. 6B. The extracellular side of the transmembrane helix was fused to an anti-ALFAtag nanobody (ALFAnb) for capturing into bNDAs obtained by stamping BSA conjugated with the ALFAtag (^ALFA^BSA). Axin 1/2 double knockout (DKO) HeLa cells co-expressing this bifunctional transmembrane crosslinker ALFAnb-TMD-GFPnb and mEGFP-Axin1 cultured on ^ALFA^BSA bNDAs showed efficient nanopatterning of Axin1 (Fig. 6C). Strikingly, the Axin1 bNDAs showed characteristic topological features of protein nanodroplet arrays, which was corroborated by DIC and SEM imaging (Fig. 6D, and Fig. S5A-D, I-L). Such features were not observed for Lrp6-based signalodots, suggesting different physicochemical properties of the nanodroplets.

**Fig. 6.**
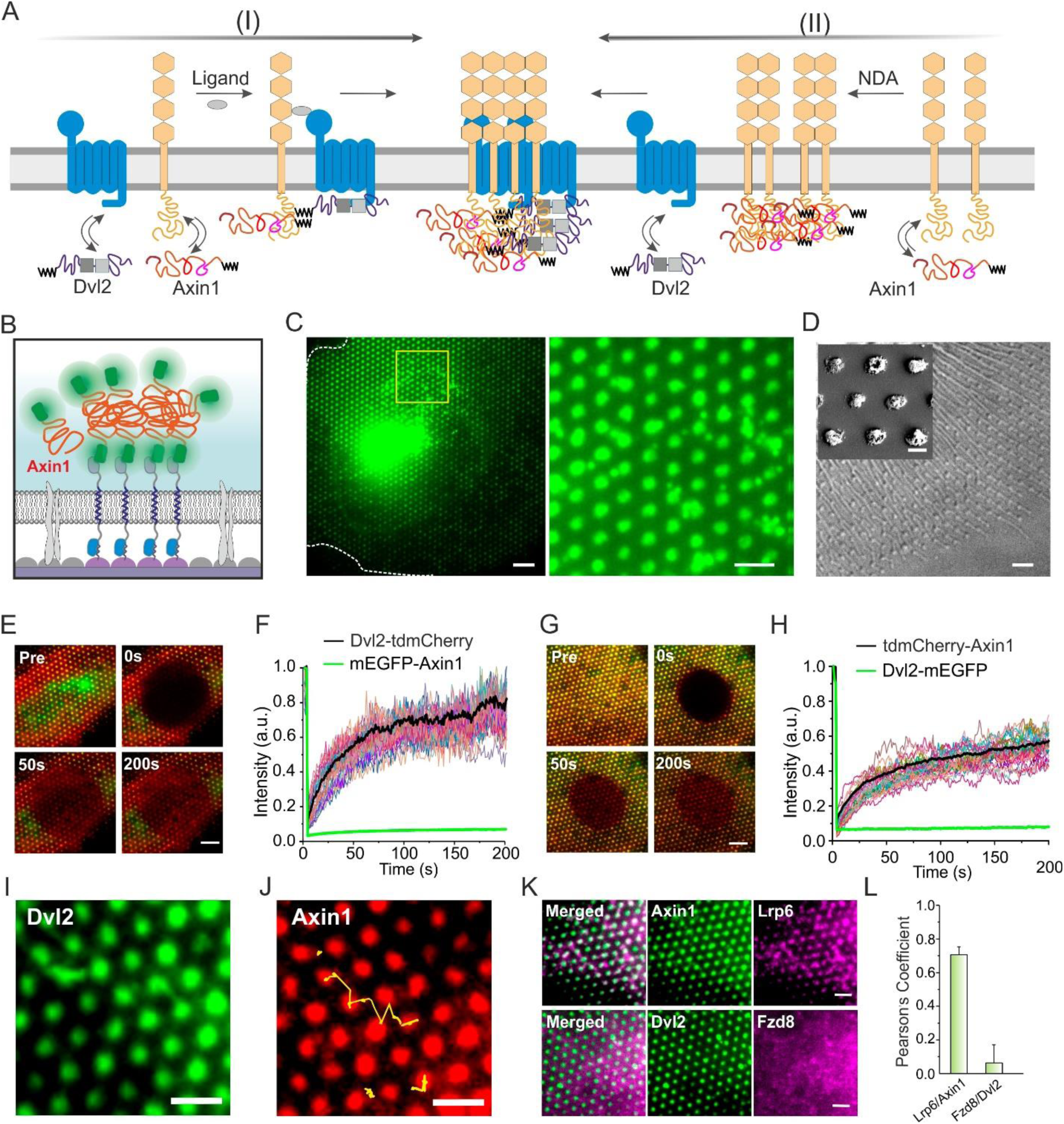
Wnt signalosome assembly driven by Axin1/Dvl2 co-condensation with co-receptors. (A) Formation of Wnt signalosomes mediated by co-condensation of Axin1 and Dvl2 can be induced by Wnt agonists that heterodimerize Fzd and Lrp6 (I) or by clustering of Lrp6 via bNDA (II). (B) Cartoon of capturing mEGFP-Axin1 into ^ALFA^BSA bNDAs using an artificial transmembrane crosslinker ALFAnb-TMD-GFPnb. (C) TIRF imaging of a HeLa cell co-expressing mEGFP-Axin1 and ALFAnb-TMD-GFPnb that were cultured on ^ALFA^BSA bNDAs. Overview (left) and zoom-up (right) of a region highlighted in yellow. Scale bar: 5 µm (2 µm in the inset). (D) Differential interference contrast (DIC) image indicates mEGFP-Axin1 bNDA being nanodroplet arrays. Scale bar: 2 µm. Inset: Scanning electron microscopy (SEM) of individual mEGFP-Axin1 nanodroplets. Scale bar: 500 nm. (E, F) Co-condensation of Dvl2 upon capturing mEGFP-Axin1 into bNDA. (E) Time-lapse dual-color fluorescence imaging. Red: Dvl2-tdmCherry. Green: mEGFP-Axin1. Scale bar: 5 µm. (F) FRAP curves obtained from 30 individual nanodroplets. (G, H) Co-condensation of Axin1 upon capturing Dvl2-mEGFP into bNDA. (G) Time-lapse dual-color fluorescence imaging. Red: tdmCherry-Axin1. Green: Dvl2-mEGFP. Scale bar: 5 µm. (H) FRAP curves of 30 individual nanodroplets. (I) Dvl2-mEGFP nanodroplets (green). (F) tdmCherry-Axin1 nanodroplets (red). Scale bars: 2µm. Yellow trajectories mark the moving Axin1 droplets. (K) Receptor recruitment by Axin1 bNDA (top) and Dvl2 (bottom) bNDA. Scale bars: 2 µm. (L) Correlation analysis of receptor recruitment into the nanodroplet arrays.

As expected, co-expression of Dvl2-tdmCherry yielded recruitment into mEGFP-Axin1 nanodroplet bNDAs (Fig. 6E), suggesting co-condensation as observed in the cytosol. FRAP experiments, however, revealed that Dvl2, but not Axin1, was rapidly exchanged in these co-condensates (Fig. 6E, F, and Fig. S6A). Likewise, capturing of Dvl2-mEGFP into bNDAs via transmembrane crosslinker yielded formation of nanodroplets (Fig. S5E-H) and co-condensation of Axin1 (Fig. 6G). FRAP of Axin1 captured into Dvl2 bNDAs showed (56 ± 7.6) % recovery within 200 s (N=30), while negligible exchange was observed for Dvl2 (Fig. 6G, H). Of note, FRAP of the artificial co-condensation of either Axin1 or Dvl2 yielded a constantly ∼40% higher recovery than that observed in Lrp6 bNDA-based signalodots (cf. Fig. 5B, D). These results highlight the different properties of Axin1/Dvl2 co-condensates depending on the subcellular context and the interactions with receptors. These differences lead to a heterogeneous intensity distribution of β- catenin immunofluorescence staining which impeded the quantification of β-catenin level.

Strikingly, we observed recruitment of Lrp6 into the artificial Axin1 bNDAs (Fig. 6K, L and Fig. S6C, G), while Dvl2 nanodroplets did not recruit Fzd8 (Fig. 6, K, L and Fig. S6D, H). These results further supports our hypothesis that membrane-proximal condensation of Axin1 initiated by Lrp6 clustering drives Wnt signalosome formation, in line with the key role of Axin that has been previously identified (59). Another unexpected observation of the Axin1/Dvl2 co-condensates is the non-correlated mobility of Dvl2 and Axin1 droplets in bNDAs (Video S7, Fig. S6E, F.). This intriguing observation suggests that two types of LLPS for Axin1 and Dvl2 co-exist within the co-condensates. The internal heterogeneity could arise from the fact that Axin1 and Dvl2 each has their unique condensation-sensitive regions to form their own droplets (60, 61) while the shared DIX homologue accounts for inducing joint LLPS (43). This result points out that hierarchical organization of LLPS occurs as a common phenomenon as observed previously in stress granules (62).

## Discussion

Spatial reorganization of cell surface receptors by micropatterning (45, 63–74) and nanopatterning (75–85) of ligands and receptors have emerged as powerful tools to unravel key determinants of collective cell functions such as adhesion, growth and differentiation as well as cellular signaling.

Spatial control of signaling platforms into nanoscale dimensions, which is relevant for numerous signaling pathways, has remained challenging. Here, we have devised capillary nanostamping for fabrication of biofunctional bNDAs with diameters of ∼300 nm. Their size is somewhat larger than that obtained by dip-/polymer-/beam pen lithography (86–88), e-beam lithography (75–78) and block copolymer micelle nanolithography (79–82), but falls in the range of DNA-directed self-assembly of multiscale patterns (73, 74, 83–85). In contrast to these aforementioned patterning approaches, capillary nanostamping involves the use of stamps penetrated by pore systems, which may act as reservoir of inks containing nanoparticles and macromolecules (89). Capillary nanostamping using the mesoporous silica stamps in this work is a robust bench-top patterning method that can be carried out manually under ambient conditions (22, 23). Moreover, capillary nanostamping can be automatized using standard instrumentation used for polymer pen lithography and it can be carried out in multi-color mode (90). The rapid, robust and cost-effective fabrication workflow makes capillary nanostamping very suitable for cell biological applications, which require high throughput and relatively large functionalized areas. The achieved dot dimensions within the diffraction limit of light are an ideal compromise for quantitative fluorescence imaging with maximum signal-to-background.

Capturing of cell surface receptors via bNDAs densely coated with a bivalent GFP clamp ensured close proximity of receptor dimers. The DARPin-based GFP clamp is a highly stable protein (53), which – in contrast to anti-GFP nanobodies – reliably maintains structure and function at the substrate surface even during cell culture at 37°C. In conjunction with high-density, high contrast capillary nanoprinting of BSA conjugates, we achieved robust capturing of proteins in the plasma membrane of live cells into densities of ∼10,000 molecules/µm². Such clustering into nanoscale dimensions with high receptor density and dimerization at molecular is highly relevant to closely mimic native signaling platforms. While requiring overexpression of the relevant receptor and effector proteins, bNDAs enable to systematically explore the correlation of receptor density and effector recruitment.

We here particularly focused on the capabilities to employ bNDAs for interrogating and controlling protein condensation at the plasma membrane, which is currently emerging as a fundamental principle responsible for dynamic assembly of signaling platforms (16). Using Wnt signalosome as a prominent example, we found that capturing Lrp6 into bNDAs promotes agonist-independent recruitment of the co-receptor Fzd8 and the cytosolic effector proteins Axin1 and Dvl2. Increased β-catenin expression levels confirmed downstream signaling activity of these Wnt signaldot arrays. Highly parallel bNDAs providing different receptor densities at single cell level enabled systematic correlation of interactions and analyzing their dynamics. These studies revealed the key role of Lrp6 density for signalodot assembly with molecular distances (∼10 nm) between Lrp6 molecules being required to efficiently initiate recruitment of all three partners. Interestingly, this threshold matches the average distance of Axin1 in co-condensates with Dvl2 we found upon co-overexpression of both proteins in the cytosol. These observations strongly support that Wnt signalosome formation is driven by co-condensation Axin1 and Dvl2 together with the receptors Lrp6 and Fzd8, which in turn is counteracting the destruction complex. Strikingly, bNDA could also be used to directly capture Axin1/Dvl2 co-condensates to the plasma membrane. Distinct morphology and phase behavior corroborate that interactions with the receptors alter the properties of Axin1/Dvl2 co-condensates, which likely is related to its functional switch. These applications highlight the unique capabilities of biofunctional bNDAs to investigate signaling-regulatory condensates at the plasma membrane of live cells, which so far have remained challenging to control (91, 92).

## Experimental Section

### Materials

Details of all used reagents can be found in the supplementary Table S3. If not mentioned otherwise, materials were used directly as purchased or received.

### Protein expression, purification and labeling

To generate tandem GFP binders used for surfaces bio-functioned with HTL, a construct of anti-GFP clamp R7 (53) followed by a (Gly-Ser-Asp)_3_ linker, the HaloTag, a second R7 and an octa- HisTag (tdClamp-HaloTag) was cloned into a pET21a vector. The tdClamp-HaloTag protein was produced in *Escherichia coli* BL21 (DE3) Rosetta cells. The cells were grown in the presence of ampicillin to an OD_600_ of 0.6 and induced with 0.5 mM IPTG at 37°C for 5 h. For purification, cells were resuspended in HBS buffer (20 mM HEPES, pH 7.5, 300 nM NaCl) supplemented with DNase, lysozyme and EDTA-free protease inhibitor mix. After disrupting the cells by sonication and ultracentrifugation, the supernatant was filtered and loaded onto a 5 mL HisTrap HP column (GE Healthcare) for immobilized metal affinity chromatography. The pooled fractions containing tdClamp-HaloTag were purified by size exclusion chromatography on a HighLoad Superdex 200 pg 16/60 column (GE Healthcare). Aliquots from combined fractions were frozen in liquid nitrogen and stored at -80°C until usage.

mEGFP with an N-terminal hexahistidine tag (mEGFP), mEGFP with an N-hexahistidine tag-HaloTag (HaloTag-mEGFP), and mCherry with an N-terminal hexahistidine tag (mCherry) were cloned in the pET21a plasmid, respectively. The proteins were expressed in *Escherichia coli* BL21 (DE 3) Rosetta cells, followed by purification of immobilized metal affinity chromatography and size exclusion chromatography as previously reported (93). The next generation surrogate (NGS) Wnt was produced as reported before (34).

Dvl2 flanked with N-terminal mEGFP and C-terminal hexahistidine tag was cloned in the pET21a plasmid and produced in *Escherichia coli* BL21 RIL cells. The cells were grown in the presence of ampicilin and chloramphenicol to an OD_600_ of 0.5. Protein expression was induced with 0.5 mM IPTG at 37°C overnight. For protein purification, cells were resuspended in a UREA buffer (8 M urea, 20 mM HEPES pH 7.5, 150 mM NaCl). The cells were disrupted by sonication and the cell lysate was subjected to ultracentrifugation. The supernatant was filtered and loaded onto a 5 mL HisTrap HP column (GE Healthcare) equilibrated in UREA buffer. After buffer exchange with HBS followed by gradient elution with HBS containing 1M imidazole, the fractions were pooled and mixed with 50 mM EDTA and 20 mM DTT for 1h. The pooled fractions were then loaded onto a Superdex 200 10/300 gel filtration column (GE Healthcare) for size exclusion chromatography in HBS buffer. Aliquots were frozen in liquid nitrogen and stored at -80°C until usage.

### Synthesis of BSA conjugates

For synthesis of ^HTL^BSA, a bovine serum albumin solution (1 g in 5 mL HBS buffer) was dialyzed in 2 L HBS buffer for 48 h at 4°C (14 kDa cutoff). Concentration of the remaining purified BSA was determined by UV-vis spectrometer. 369 µL the dialysis-purified BSA (0.37 µmol) was added to 3.4 mL HBS together with a solution of 96 mg EDC (0.5 mmol) in 1 mL HBS. 1.7 mg HTL-NH_2_ (7.6 µmol) in 200 µL ethanol was then added to the reaction mixture and shaken overnight at 4°C. The reaction mixture was dialyzed in 2 L HBS for 24 h at 4°C (14 kDa cutoff), resulting in a 62.8 µM ^HTL^BSA solution. MALDI analysis showed a broad peak at 69334 m/z compared to the unfunctionalized BSA (66294 Da measured by MALDI). The average mass difference of 3081 m/z corresponds to a degree of functionalization ≈ 13.6 HTL moieties per BSA molecule.

For synthesis of ^ALFA^BSA, 4 mg cysteine-containing ALFA peptide (Ac-CPSRLEEELRRRLTE- COOH, 2.0 µmol) was dissolved in 300 µl HBS (pH: 6.5) followed with addition of 1.3 mg Mal-PEG_2_-NHS (3.1 µmol). The mixture was shaken for 15 min before adding 259.2 µl of the dialysis purified BSA (0.41 µmol). The solution was gently shaken over night at 4°C and dialyzed in HBS for additional 48 h at 4°C (14 kDa cutoff), resulting in a 151 µM ^ALFA^BSA solution. MALDI analysis showed convoluted broad peaks at 66932 m/z, 69272 m/z and 71267 m/z indicating a mixture of un-functionalized BSA, mono- and bi-functionalized ^ALFA^BSA with an estimated molar ratio of 1:1.2:1.

^AT647N^BSA was synthesized by mixing the purified BSA (200 µL, 1.5mM) and ATTO647N-NHS in a 1:1 molar ratio in HBS buffer for 4h at room temperature. The crude product was purified with an Amicon Ultra (3K) centrifugal filter and the supernatant was used for stamping.

### Surface modification on glass

Glass substrates and silica transducers for reflectance interference measurements (the RIFs chips) were cleaned with a lint-free cloth and plasma cleaned for 10 min in a Diener Femto plasma cleaner prior to the silanization. Immediately after cleaning, the substrates were immersed into a 2% v/v silane solution in dry toluene for 1 h. After the given time the substrates were rinsed thoroughly with toluene and ethanol to remove residual unreacted silane and used within a timeframe of 2 h. This procedure was applied for all silanes with the exception of octadecyltrichlorosilane, which was taken out after a reaction time of 10 min to avoid un-controlled polymerization.

### Solid phase detection by TIRFS-RIF

A home-built set-up for total internal reflection fluorescence spectroscopy (TIRFS) and reflectance interference (RIF) was used as described in detail before (49, 94). Surface sensitive detection of protein binding kinetics were determined in the TIRFS-RIF setup under flow conditions. For instance, for TIRFS-RIF detection of ^HTL^BSA on an NMTS surface, 10 µM ^HTL^BSA in HBS buffer (20 mM HEPES pH 7.5, 150 mM NaCl) was injected into the flow chamber. Binding of HaloTag- mEGFP to ^HTL^BSA-covered NMTS surfaces was measured by injecting 2 µM of HaloTag-mEGFP and tagless-mEGFP (negative control) separately into the system. Mass signals and fluorescence signals were recorded simultaneously in RIF channel and TIRFS channel, respectively, which were used for binding kinetic analysis of protein binding (BIAevaluation 3.0 software).

### Capillary nanostamping

Mesoporous silica stamps were prepared following procedures described by Schmidt et al. (23). Water-soaked stamps were washed by placing them in ethanol for approximately 1 day and refreshing the ethanol at least three times. The ethanol-soaked stamps were immersed into MilliQ water which was repeatedly refreshed until the solvent was exchanged. Then the stamps were infiltrated for ≈ 30 min with 15 µM ^HTL^BSA or ^ALFA^BSA in HBS. In order to remove the excess ink, the stamp was dried under a stream of nitrogen until a clear rainbow shine was visible. After attachment to the stamp holder via double-sided tape, the NMTS modified glass substrates were subjected for patterning for ∼3-5 s and backfilled with 100 % fetal bovine serum (FBS). For multiple patterning cycles, new inks were added on the structured stamp surface for ≈ 1 min and blown dry afterwards. Stamping with fluorescently labelled BSA was performed by using 1.6 µM ^AT647N^BSA in HBS as ink. The patterned samples were backfilled with FBS for 10 min and washed with HBS. The patterned slides were washed with water and stored in PBS until further use.

### Cell culture, transfection & immunofluorescence

HeLa cells (DSMZ no.: ACC57, German Collection of Microorganisms and Cell Cultures) were cultivated at 37°C, 5 % CO_2_ in Dulbecco’s Modified Eagle’s Medium supplemented with 10% FBS. Axin1/2 DKO HeLa cell line was generated by Cyagen US Inc. via CRISPR/Cas9 gene knockout (Contract No. SCS181212JM1). The gRNAs used for Axin1 are: (1) AGACTTCGACGGCCACTCCG<u>AGG</u> and (2) TCCAGTAGACGGTACAGCGA<u>AGG</u>. gRNAs for Axin2 are: (1) ACACCAGGCGGAACGAAGAT<u>GGG</u> and (2) GTGCAAACTTTCGCCAACCG<u>TGG</u>.

A homozygous clone was selected out. Double knock-out of Axin1 and Axin2 in the homozygous clone was validated by RT-PCR (Axin1: FW-primer: CTCTGGTCGTGTTTCATGGACC. BW- primer: CCTGAAACGTCCACTCCTCC; Axin2: FW-primer: GCCGATTGCTGAGAGGAACT. BW-primer: AGTCAGCACTGGAAGACTGC) and DNA sequencing. The results showed genetical deletions of 756bp for Axin1 and 563 bp for Axin2 at Exon 2, respectively. Due to the deletions, the residual Axin1 contains 58 aa of the original N-terminal Axin1 and stops at 114 aa (hAxin1 full length 862 aa). The residual Axin2 contains 58aa of the N-terminal and stops at 79 aa (hAxin2 full length 843 aa).

Transient transfection was carried out by a polyethyleneimine (PEI) transfection protocol. Details of protein plasmids can be found in the Supplementary Table S2. Briefly, HeLa cells were plated onto a 6 cm Petri dish and grown overnight. 5 µg of each plasmid was added to 300 µL 150mM NaCl, then vortexed and mixed with 10 µL PEI. After 15 min incubation at room temperature, the mixture was transferred to the seeded HeLa cells for 2-3 h, followed with twice PBS washing and kept in the incubator overnight. For experiments of artificial LLPS, transfections of HaloTag-Axin1 and Dvl2-tdmCherry were performed according Viafect^TM^ protocol (Promega) with a ratio of 1 µg DNA to 3 µl Viafect^TM^ reagent and kept overnight in incubator. The transfected HeLa cells were detached from the petri dish by Trypsin/EDTA treatment and seeded on a prepared coverslip. After incubation of 6-15 h, the samples were used for microscopy imaging. For labelling of Fzd8- SNAP on bNDA, 100 nM SNAP-Surface 549 or SNAP-Surface 647 was added to the cell culture medium for 30 min at 37 °C. Labeling of HaloTag was conducted by adding 50 nM SiR-HTL for 20 min at 37 °C. For single-molecule tracking experiment, 500 pM SiR-HTL was used for sub- stoichiometric labeling of HaloTag-Dvl2. After labeling, the cells were washed trice with PBS and then immersed in cell culture medium for microscopy imaging experiments.

For immunostaining, cells were fixed with 1 mL 4% PFA in HBS for 15 min at RT. Afterwards, the fixation mixture was slowly siphoned off and the glasses were washed with PBS three times for 5 min each. The fixed cells were subjected to 0.1 % v/v solution of Triton-X100 in PBS for 10 min, washed three times with PBS and blocked with a solution of 3 % m/m BSA in PBS for 30 min. After removing the blocking solution, 20 µL of a 1:1000 diluted antibody solution with 3 % BSA was added to the coverslips and topped off with a small piece of parafilm. The sample was placed in a loosely closed container and put into the 4 °C fridge overnight. The samples were washed three times again with PBS and placed into a microscopy sample holder for fluorescence imaging.

### Total internal reflection fluorescence microscopy (TIRFM)

TIRF imaging was carried out at 25 °C using an inverted microscope (Olympus IX-83) with two decks equipped with a 4-Line TIRF condenser (cellTIRF (MITICO), Olympus) in the upper deck and a cellFRAP module (Olympus) in the lower deck. The setup was equipped with a 405 nm laser (BCL-100-405, CrystaLaser), 488 nm diode laser (LuxX 488-200, Omicron), as well as a 561 nm (2RU-VFL-P-500-561-B1R, MPB Communications) and 642 nm fiber laser (2RU-VFL-P- 500-642-B1R, MPB Communications). Fluorescence excitation and detection was accomplished by using a TIRF pentaband polychroic beamsplitter (Semrock zt405/488/561/640/730rpc) and a penta-bandpass emitter filter (BrightLine HC 440/521/607/694/809). Emission was further filtered by single bandpass filters used for each channel (blue: Semrock BrightLine HC 445/45, green: Semrock BrightLine HC 525/35, orange: Chroma 600/50 ET, red: Chroma 685/50). Images were acquired with an electron-multiplying CCD camera (Andor iXon Ultra 897) using either a 150x oil immersion objective (Olympus UAPON 150× TIRF, NA 1.45) or a 100x oil immersion objective (Olympus UPLAPO 100× HR, NA 1.5) with and without an additional 1.6x magnification (IX3-CAS, Olympus). Differential interference contrast (DIC) images were taken with the standard DIC condenser of the microscope. The software for imaging was CellSens Dimension (Version 3.1, Olympus).

For fluorescence recovery after photobleaching (FRAP) on bNDAs, the Olympus cellFRAP module was used in combination with a 405 nm bleaching laser (LuxX+ 405-60, Omicron). Images were acquired with a 3-Line polychroic beamsplitter (Chroma, zt488/561/633,405tpc) in the upper deck combined with a FRAP cube in the lower deck consisting of a dichroic beamsplitter at 405 nm (Chroma, H 405 LPXR superflat) with a QuadLine emitter filter (Chroma, zet405/488/561/640m). Three pre-bleaching images were recorded for a circular region of interest (ROI) with a ∼20 µm diameters at full laser power. Images were acquired at the time interval of one frame per second for 4 min. Within the bleached area, individual nanodots (N=20-40) were selected and the fluorescence intensity of each was measured separately (*F*_*nanodot*_(*t*)). The intensities of each nanodot *F*(*t*) were corrected against photobleaching using the following equation:

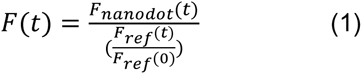

With *F*_*ref*_ (*t*) being the fluorescence of unbleached dots and *F*_*ref*_ (0) the intensity of the same dots before bleaching.

Direct stochastic optical reconstruction microscopy (dSTORM) image sequences were acquired with 500 µl of a 2-amino-2-(hydroxymethyl)propane-1,3-diol (TRIS) imaging buffer (150 mM TRIS pH 8.5, 10 mM NaCl) containing 2 % β-mercaptoethanol, 10 mg/ml glucose, 0.04 mg/ml catalase, 0.5 mg/ml glucose oxidase as an oxygen scavenger system. To achieve stable single-molecule blinking conditions, the sample area was illuminated with the 642 nm laser at full intensity for roughly 2 min to transfer most AT647N to into the dark state. Raw data sets consisted of 30.000 frames acquired at 40 fps with an EM gain of 5 at a pixel size of 160 nm. To enable reactivation from the long lived dark state, an additional constant low power UV illumination at 405 nm was used. Sample focus plane was held during acquisition with a piezo z-stage (NanoScanZ, NZ100, Prior Scientific). Raw datasets were further processed with the ImageJ plugin ThunderSTORM (95). Parameters for image processing were as follows: Image filtering was done via a wavelet (B-spline) filter with order 3 and scale 2, approximate localization of molecules was conducted via a local maximum estimation with a peak intensity threshold equaling 2 times the standard deviation of the wavelet filter used for signal processing. Sub-pixel localization of molecules was performed with an integrated Gaussian method using a fitting radius of 5 pixels and weighted least squares fit.

### Single molecule tracking

For single molecule tracking in Dvl2 condensate, HaloTag-Dvl2 and mCherry-Dvl2 were co-expressed in HeLa cells using a pSems vector (Table S2). Sub-stoichiometric labeling of HaloTag-Dvl2 with 500 pM of SiR-HTL was used. Dual-color TIRF microscopy imaging by 561nm and 642nm excitation was carried out in the above-mentioned Olympus TIRF 4-Line setup. Simultaneously time-lapse imaging of the 561 and 642 channels was acquired with a frame rate of 30 fps. To highlight the Dvl2 condensation, images of the mCherry-Dvl2 channel (561 channel) were Z-projected into an overlapping image using Image J software (v1.49i)(96). Single molecule tracking and the step length histogram analysis of the SiR-labeled HaloTag-Dvl2 was performed as previously reported (97, 98). A triple-component Gaussian model was found to be the optimal fitting model for quantifying the heterogenous diffusion of Dvl2 in the condensates.

### Confocal laser scanning microscopy (cLSM)

cLSM imaging was carried out on a confocal laser-scanning microscope (FluoView 1000, Olympus) equipped with a 60× water-immersion objective (UPLSAPO, NA1.2, Olympus). The microscope was equipped with different filter cubes for the orange (U-MWG2, Olympus) and green channel (Exciter bandpass: 474-491 nm, Semrock BrightLine HC 482/18; Emitter bandpass: 505-545 nm, D525/40; Beam splitter: 488 nm and 647 nm, HC-DB 488/647). Concentration calibrations curves were obtained by using purified mCherry at different gains. For FRAP experiments in cLSM, a circular ROI was selected (8 μm diameter for whole droplet FRAP and 5 μm for sub-droplet FRAP). Three pre-bleach images were recorded before the ROI was illuminated by 405 nm. A time zero intensity *I_0_* was recorded from the center of the ROI immediately after photobleaching. Fluorescence recovery was monitored with a 4.05 s time interval. The fluorescence intensity in the ROI was obtained by subtracting the time-zero intensity, i.e. (*I*_*ROI*_ (*t*) − *I*_0_). A reference region outside the photobleached ROI was processed in the same way to obtain *I*_*REF*_ (*t*). The ROI intensity was normalized to the reference for each time interval using Equation (2):

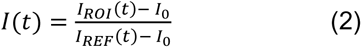

where *I_0_* is the time-zero intensity after photobleaching, *I*_*ROI*_ (*t*) is the fluorescence intensity within the ROI, and *I*_*REF*_ (*t*) is the intensity outside the photobleaching ROI. *I*(*t*) is the time-dependent intensity used for plotting the FRAP curve.

### Scanning electron microscopy

Cells for scanning electron microscopy (SEM) were fixed 1 mL 4% PFA in HBS and 1% glutaraldehyde for 1 h at RT. The samples were washed with PBS buffer trice and transferred into an ethanol solution by slowly increasing the ethanol content. Critical point drying was conducted with a Leica Auto CPD300 with liquid CO2 at 40 °C and 80 bar. The samples were sputtered with platinum/iridium before imaging with a Zeiss Auriga scanning electron microscope operated at an accelerating voltage of 3 keV. Both secondary electron secondary ion detector (SESI) and in-lens detector were used for detection.

### Image evaluation

Immunostaining of β-catenin: For every cell, the nucleus was determined through the DIC image and the contour was transferred to the respective fluorescence image in order to determine the intensity values inside the nucleus. Images were acquired under the same conditions for each set of experiments. The results of each set was referenced against untreated wild type HeLa cells for reliable comparison.

Contrast analysis: Contrast analysis of line profiles was performed based on deconvolved images. Values from a line plot were subtracted either with a background outside a cell, when working *in cellulo,* or with a defective spot in the pattern to minimize the effects of intensity overlapping of two neighbor dots. Box-chart contrast analysis of bNDA in the deconvolved image was obtained using the colocalization algorithm (cf. Colocalization analysis of bNDAs). Box-chart diagrams were gained by plotting the determined contrast of individual nanodots for each channel.

Pearsońs Coefficient: For the determination of the Pearsońs coefficient, three representative cells of the respective experiment were analyzed. For each cell, 3 line plots were drawn through the whole patterned cell and the intensity values in every fluorescence channel were recorded. Pearsońs correlation analysis was performed with the obtained values of the corresponding channels. Data are shown as mean value ± standard deviation.

Image evaluations were performed with the ImageJ (v1.49i). Origin 9 Pro was used for contrast evaluation and statistical analysis.

### Colocalization analysis of bNDAs

The colocalization script was written in Matlab R2018a containing the following three parts:

1. Image deconvolution: A Tikhonov deconvolution was performed to the raw image by reversing the effect of the point spread function (PSF) and shifting all signal back into the nanodots:

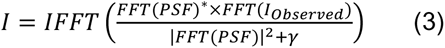

Here, PSF=1.3 pixel was used for blue, 1.4 pixel for green, 1.8 pixel for red. PSF of the microscope was approximated by 2D-Gaussian. *γ* = 10^−1.5^ was used as the regularization factor for smoothness during the deconvolution to suppress the noise enhancement.
2. Nanodot identification: Individual nanodots were identified by smoothing the deconvolved image and then seeking the local maxima in the pixel intensities. In order to discriminate signal against noise, a minimum signal threshold is estimated using Otsu’s method (99). The detected nanodots were further refined by first calculating the median distance between nanodots and then discarding all nanodots that do not satisfy at least 3 such neighbors (<1.3x median distance). The accepted pixels were then thickened radially by 2 pixels (pixel size 130 nm) to capture the whole printed area. Of note, the thickening was done in a way that there is always 1 pixel separating between individual nanodots. Additionally, a cell mask was generated by dilating the image further with 2 pixels or by filling holes if necessary.
3. Contrast calculation and classification: The contrast was calculated by comparing the intensity within a nanodot to the local background determined from the surrounding pixels. After smoothing by lateral interpolation, the average intensities in the individual nanodots and the background were calculated. The contrast was then obtained as the ratio of intensity in the nanodot divided by the background for all channels. Contrast classification was carried out to determine the minimum contrast of Lrp6 (i.e. the threshold) that induces co-recruitment of Fzd8, Axin1 and/or Dvl2. To do so, nanodot contrasts of each channel were analyzed by fitting a 2-Fraction Gaussian Mixture against the contrasts obtained in the Lrp6 channel. By setting a minimum value (here using a contrast of 2), we determined the posterior probabilities that belong to the effective response fraction (marked in red in the figures). The minimum value was then sampled via bootstrapping the 5% percentile of the Lrp6 contrast that is responsible for recruitment of proteins in the other two channels. The minimum contrast of Lrp6 was determined when the responses being close to log(ratio) = 0 by checking the contrast distributions of other channels. To this end, the power law dependencies as well as a correlation coefficient were determined together with the minimum contrast of Lrp6.

### Statistical Analysis

Data were presented as mean ± SD. For immunofluorescence staining of β-catenin, n >1 5 cells. For Pearson’s correlation analysis, n = 3 cells with three lines each. For single nanodot intensity analysis, n ≥ 700. For FRAP analysis, n ≥ 25. The Student’s t-test was used to compare the means between two independent sample groups. Statistical significance was defined by p-values of *p ≤ 0.05, **p ≤ 0.01, and ***p ≤ 0.001. Image evaluations were performed with the ImageJ (v1.49i). Origin 9 Pro was used for contrast evaluation and statistical analysis. Single nanodot intensity analyses were performed with Matlab R2018a.

### Data and materials availability

All data needed to evaluate the conclusions in the paper are present in the paper and/or the Supplementary Materials. Additional data related to this paper may be requested from the authors.

## Acknowledgement

We thank H. Kenneweg, A. Budke-Gieseking, G. Hikade and W. Kohl for the excellent technical assistance. This project was supported by funding to C.Y., J.P., and R.K. from the DFG (YO 166/1-1, SFB 944, projects P8 and Z, and the DFG Facility iBiOs, PI 405/14-1), funding to M.S. from the European Research Council (ERC-CoG-2014, Project 646742 INCANA) and by intramural funding by Osnabrück University within the profile line ”Integrated Science”.

## Author Contributions

C.Y. and J.P. conceived the project together with M.S. and M.P. M.P. designed and conducted the bNDA experiments and performed the data analysis. M.K. performed the cytosolic LLPS experiments and analyzed the results. C.P.R conceived and programmed the evaluation software for image analysis. A.P. contributed the GFP trap technology. I.W. performed the protein engineering and characterizing the binding kinetics. L.J. and S.K. contributed to the synthesis of surface coating materials and surface sensitive detection. Y.M. produced the Wnt agonist under the supervision of K.C.G. M.H. and R.K. contributed to superresolution imaging and image evaluations. C.Y. and J.P. wrote the manuscript with contributions from M.S., M.P. and all authors.

## Competing Interests

The authors declare that they have no competing interest.

## SUPPLEMENTARY FIGURES

**Fig. S1.**
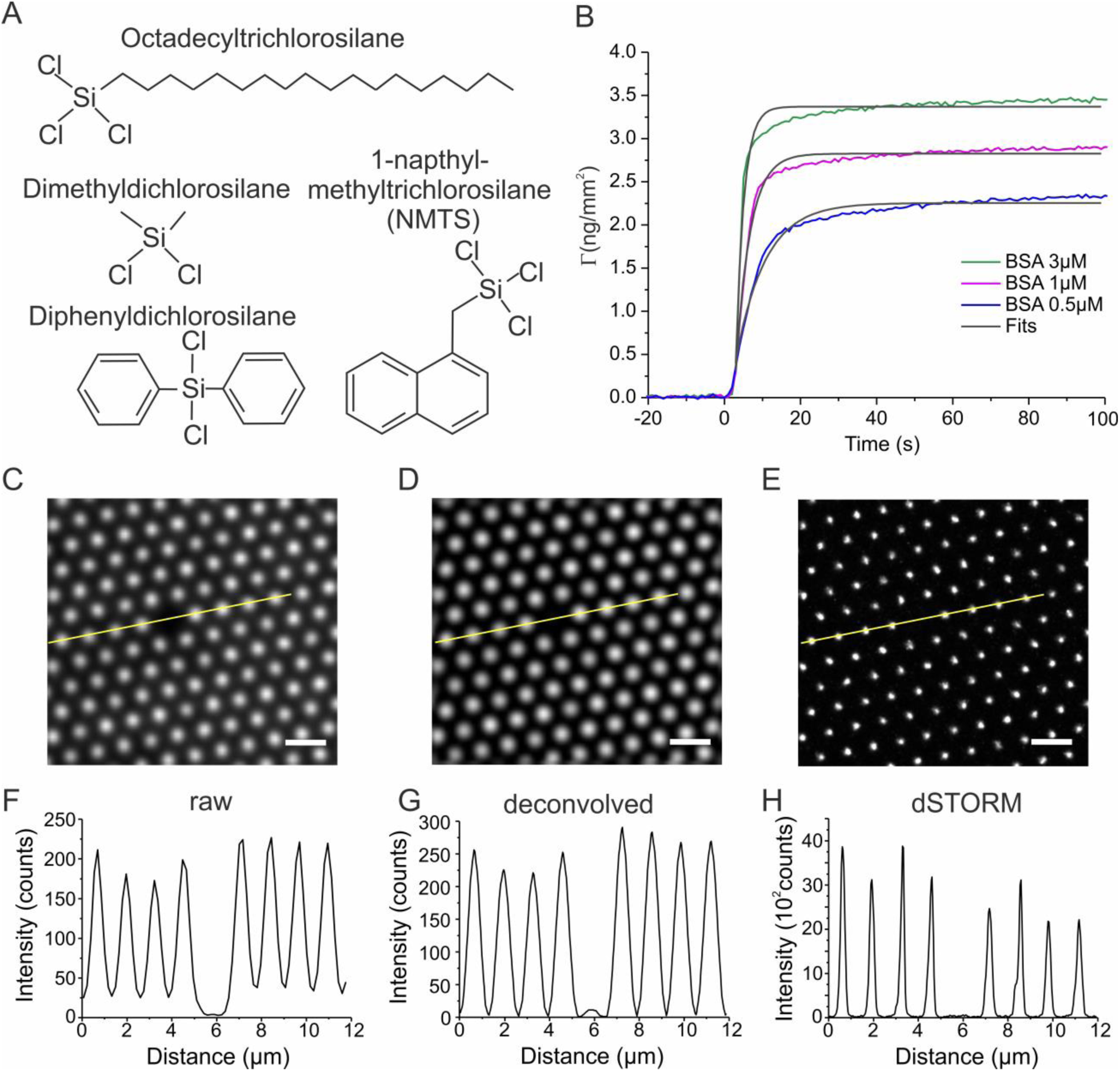
Surface modification for high-contrast protein nanopatterning by capillary nanostamping. (A) Chemical structures of silanes used for rendering silica substrates hydrophobic. (A) Binding of BSA on NMTS-coated silica monitored in real-time by reflectance interference (RIF) detection under flow-through conditions. An association rate constant *k_a_* = 2.7×10^5^ M^-1^s^-1^ was estimated from the fit (Table S1). (C-H) Signal and contrast of representative nanodot arrays (NDAs) obtained by printing ^AT647N^BSA assessed by fluorescence microscopy. Raw TIRF microscopy image (C), Deconvolved TIRF microscopy image (D) and dSTORM image (E). Scale bar 2 µm. Corresponding intensity profiles highlighted in the yellow line in the corresponding images for the raw (F), the deconvolved (G) and the dSTORM (H) images. The true background can be estimated in the defect region.

**Fig. S2.**
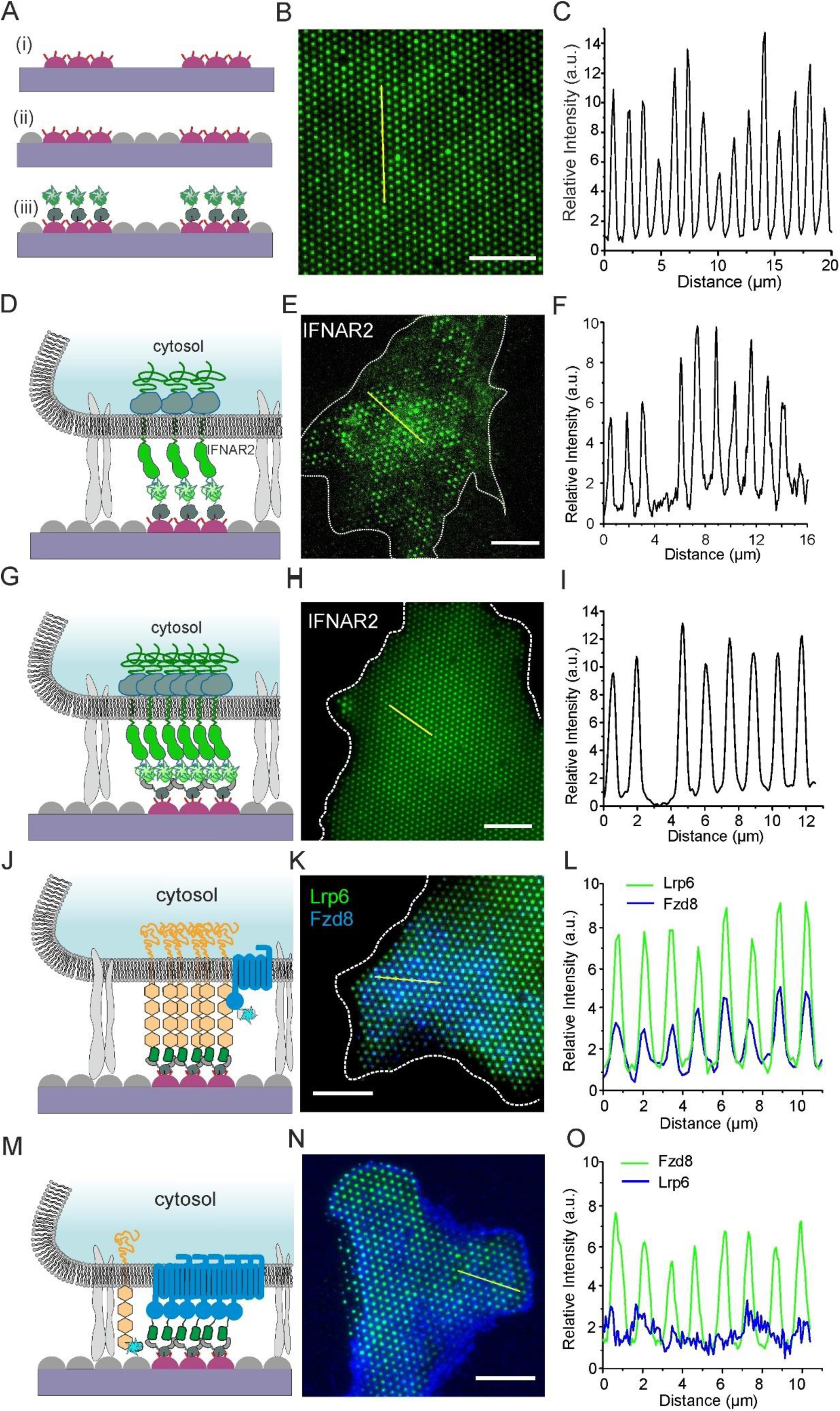
Capturing strategies into bNDAs via HaloTag and tdClamp *in vitro* and *in cellulo*. (A-C) Fluorescence staining of ^HTL^BSA bNDAs by reaction with HaloTag-mEGFP. (A) Cartoon of the surface architecture with ^HTL^BSA dots (red), FCS backfilling (grey) and HaloTag-mEGFP (green). (B) Representative TIRF microscopy image. Scale bar 10 µm. (C) Intensity profile along the yellow line highlighted in panel A. (D-F) Formation of Type-I inteferon receptor bNDAs in live cells via ^HTL^BSA functionalization. (D) Cartoon of capturing HaloTag-mEGFP-IFNAR2 on ^HTL^BSA bNDAs. (E) TIRF microscopy image of HaloTag-mEGFP-IFNAR2 bNDAs in live cell. (F) Intensity profile along the yellow line highlighted in panel E. (G-I) Formation of Type-I inteferon receptor bNDAs via tdClamp functionalization. (G) Cartoon of capturing HaloTag-mEGFP-IFNAR2 on tdClamp functionalized ^HTL^BSA bNDAs. (H) TIRF microscopy image of HaloTag-mEGFP-IFNAR2 bNDAs in live cell. (I) Intensity profile along the yellow line highlighted in panel H. (J-O) Formation of Lrp6 and Fzd8 nanopatterns in live cells on tdClamp-functionalized ^HTL^BSA bNDA. (J) Cartoon of capturing mEGFP-Lrp6 by tdClamp and co-recruitment of SNAP-Fzd8 (labeled with ^Dy647^). (K) Merged TIRF microscopy image of the immobilized mEGFP-Lrp6 (green) on tdClamp functionalized ^HTL^BSA bNDAs with the co-recruited SNAP-Fzd8 receptor (labeled with ^Dy647^, blue). (L) Intensity profile along the yellow line highlighted in panel K. (M) Cartoon of capturing mEGFP- Fzd8 to tdClamp bNDAs in presence of co-expressed mScarlet-Lrp6. (N) Merged TIRF microscopy image of mEGFP-Fzd8 NDAs (green) and mScarlet-Lrp6 (blue). (O) Intensity profile along the yellow line highlighted in panel N. Scale bars: 10 µm.

**Fig. S3.**
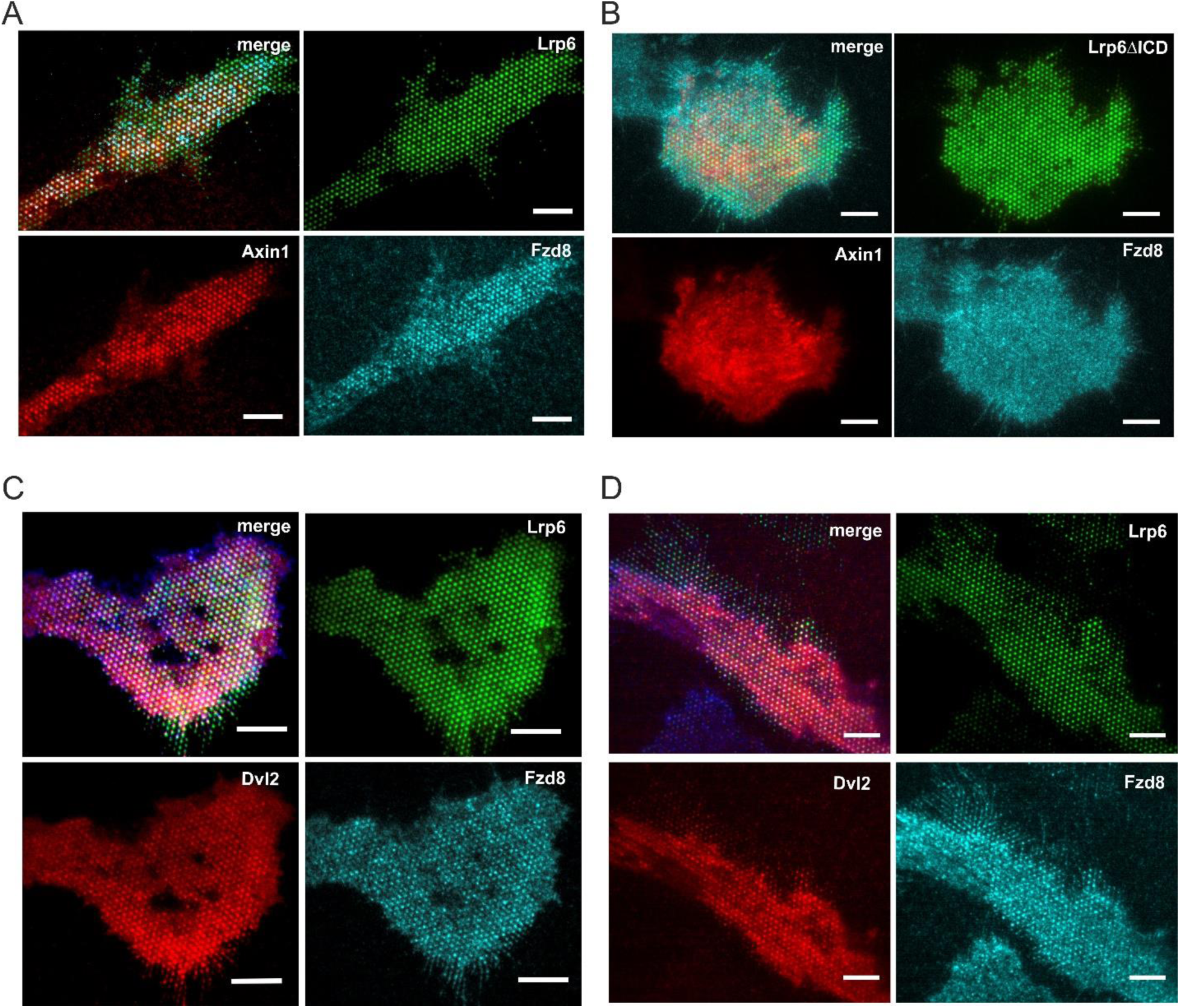
Whole cell images of co-patterning experiments on tdClamp-functionalized bNDAs. (A) TIRF microscopy image of a wild type (wt) HeLa co-expressing mEGFP-Lrp6 (green), SNAP- Fzd8 labeled with ^Dy647^ (cyan) and tdmCherry-Axin1 (red). (B) TIRF microscopy images of a wild type HeLa cell co-expressing mEGFP-Lrp6ΔICD (green), SNAP-Fzd8 labeled with ^Dy647^ (blue) and tdmCherry-Axin1 (red). (C) TIRF microscopy images of a wild type HeLa cell co-expressing mEGFP-Lrp6 (green), SNAP-Fzd8 labeled with ^Dy647^ (blue) and tdmCherry-Dvl2 (red). (D) TIRF microscopy images of an Axin1/2 double-knock-out HeLa cell co-expressing mEGFP-Lrp6 (green), SNAP-Fzd8 labeled with ^Dy647^ (blue) and Dvl2-tdmCherry (red). Scale bars: 10 µm.

**Fig. S4.**
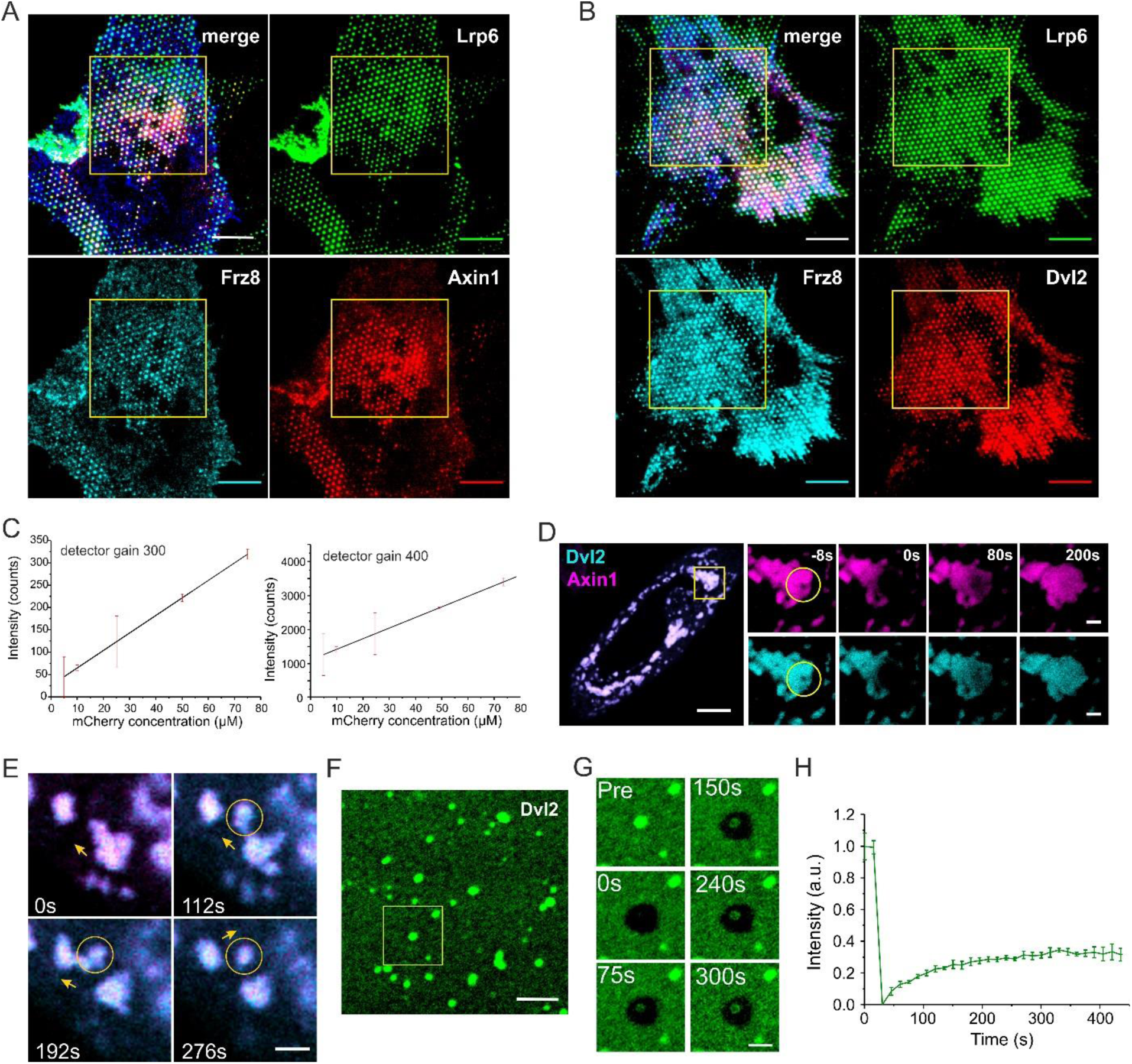
Dynamics of Axin1 and Dvl2 (co-)condensates. (A, B) Representative full-cell images of Wnt signalodots for FRAP experiments. (A) TIRF images of a wt HeLa cell co-expressing mEGFP-Lrp6 (green), SNAP-Fzd8 labeled with Dy647 (blue) and tdmCherry-mAxin1 (red) prior to FRAP. Scale bars: 10 µm. (B) TIRF images of a HeLa cell co-expressing mEGFP-Lrp6 (green), SNAP-Fzd8 labeled with Dy647 (blue) and Dvl2-tdmCherry (red) prior to FRAP. Scale bars: 10 µm. (C) Calibration plots of intensity-concentration used for quantifying the Dvl2-mCherry concentrations *in cellulo*. Purified mCherry was used in the confocal microscopy measurement with different detector gains. (D) Sub-droplet dynamics of the co-condensation of Axin1 (magenta) and Dvl2 (cyan) characterized by confocal fluorescence microscopy. Scale bar: 2 µm (10 µm in the overview image). (E) Fission and fusion of Axin1/Dvl2 co-condensates captured in time-lapse confocal microscopy (cf. Video S5). (F-H) Dynamics of Dvl2 droplets formed from purified protein in *vitro*. (F) Confocal laser scanning microscopy of 2 µM mEGFP-Dvl2 in HBS buffer. Protein droplets formation was initiated by adding 5% w/w PEG (Mw 3000). Scale bar: 5 µm. (G) Time-lapse microscopy imaging of the photobleached region marked in yellow in panel F. Scale bar 2 µm. (H) FRAP of mEGFP-Dvl2 droplets. Mean±s.d. were obtained from triple experiments.

**Fig. S5.**
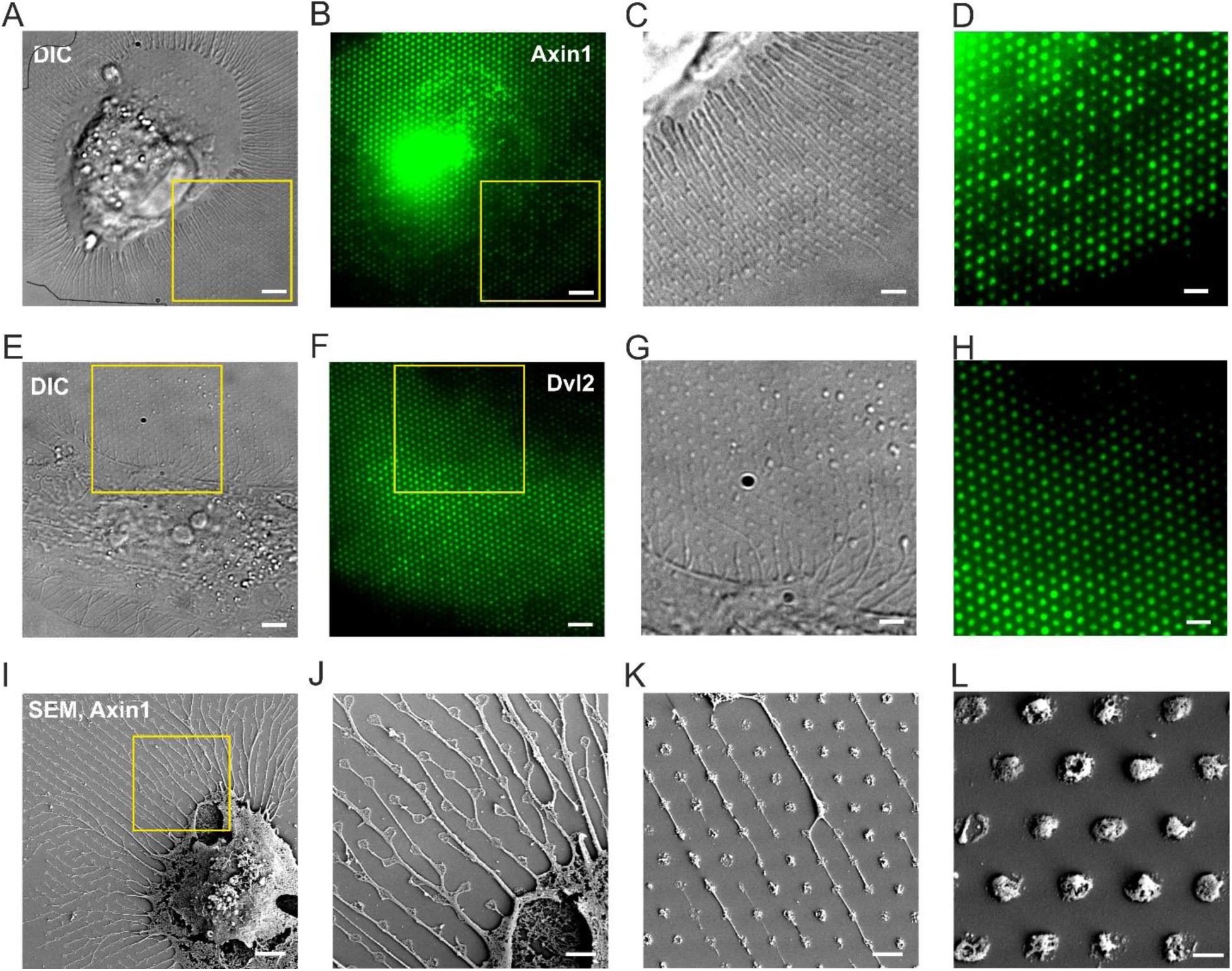
Intracellular capturing of Axin1 or Dvl2 into bNDAs using artificial transmembrane anchors ALFAnb-TMD-GFPnb. (A-D) Capturing of Axin1 on bNDAs in HeLa cells. (A) Differential interference contrast (DIC) image of a HeLa cell co-expressing mEGFP-Axin1/SNAP-Fzd8 showing distinct mEGFP-Axin1 nanodroplet arrays. (B) TIRF microscopy image of mEGFP-Axin1 in the cell. Scale bars: 5 µm. (C, D) Zoom-up DIC image (C) and TIRF microscopy image (D) of the highlighted area. Scale bars: 2 µm. (E-H) Formation of Dvl2 bNDAs using artificial transmembrane anchors. (E) DIC image of a HeLa cell co-expressing mEGFP-Dvl2/SNAP-Fzd8 showing mEGFP-Dvl2 phase separation in nanodroplets arrays. (F) TIRF microscopy image of mEGFP-Dvl2 in the cell. Scale bars: 5 µm. (G, H) Zoom-up DIC image (G) and TIRF microscopy image (H) of the highlighted area in (E). Scale bars: 2 µm. (I-L) Scanning electron microscopy (SEM) images of a mEGFP-Axin1/SNAP-Fzd8 co-expressing HeLa cell on bNDAs. (I) SEM of a fixed HeLa cell with mEGFP-Axin1 nanodroplet arrays. Scale bar: 3 µm. (J) Zoom-up image of the highlighted area in (I). Scale bar: 1 µm. (K) Another representative SEM image of mEGFP- Axin1 bNDAs. Scale bar: 1 µm. (L) SEM image of individual mEGFP-Axin1 nanodroplets. Scale bar: 500 nm.

**Fig. S6.**
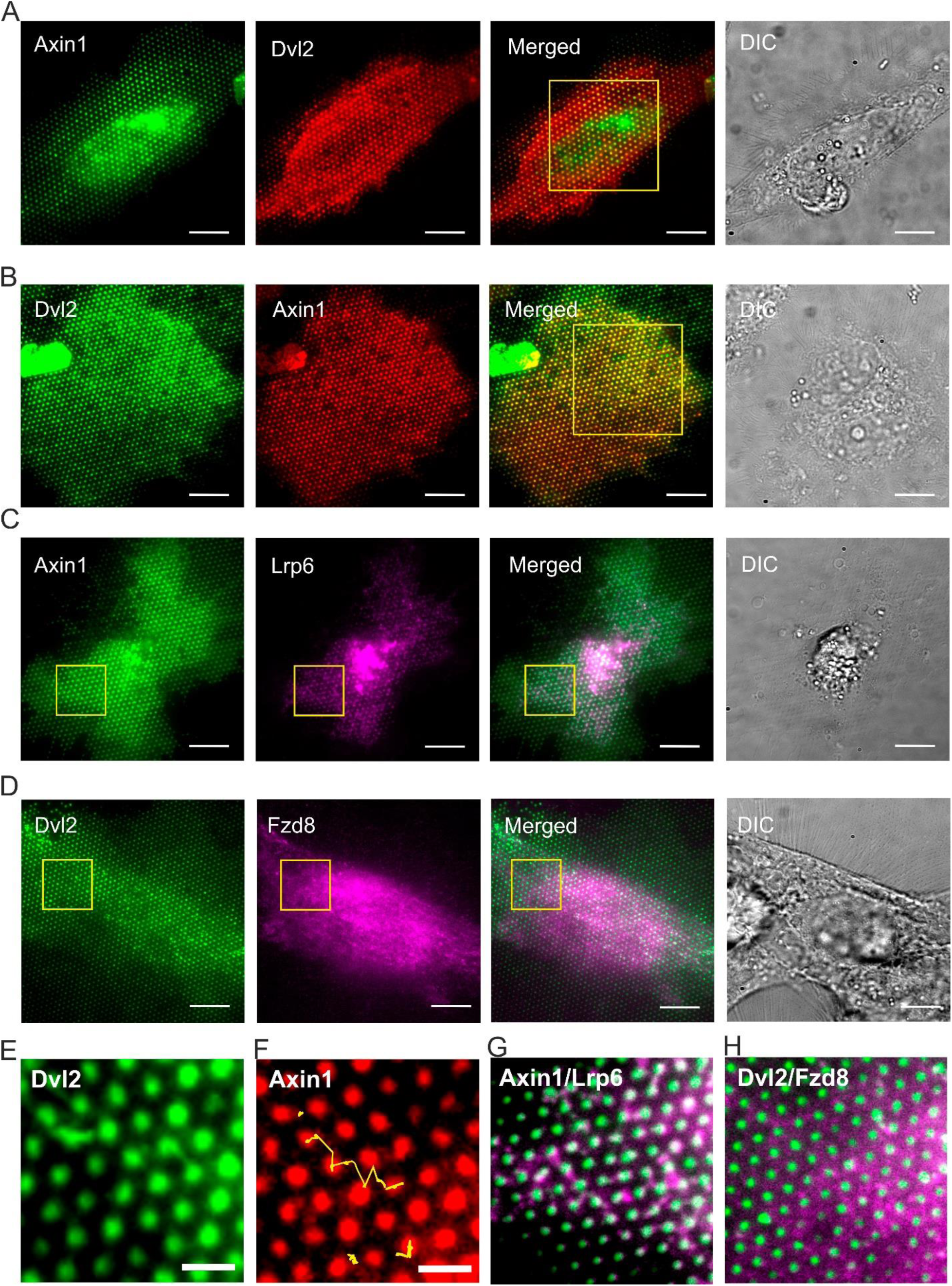
Condensations of Axin1 and Dvl2 and their interactions with co-receptors in Axin DKO HeLa cell. (A) Dual-color TIRF microscopy images of mEGFP-Axin1 bNDA (green) and Dvl2- tdmCherry (red). (B) Dual-color TIRF microscopy images of mEGFP-Dvl2 bNDA (green) and tdmCherry-Axin1 (red). Scale bars: 10 µm. Yellow box marks the FRAP region. (C) Dual-color TIRF microscopy images of mEGFP-Axin1 bNDA (green) and mScarlet-Lrp6 (magenta). (D) Dual-color TIRF microscopy images of mEGFP-Axin1 bNDA (green) and mScarlet-Fzd8 (magenta). Scale bars: 10 µm. Yellow boxes mark region for correlation analysis. (E, F) Intensity projection of 240 time-lapse TIRF microscopy images for Dvl2 and Axin1 nanodroplets. Image acquisition rate: 1fps. (E) Dvl2-mEGFP nanodroplets (green). (F) tdmCherry-Axin1 nanodroplets (red). Scale bars: 2µm. Droplet dynamic of Axin1 between bNDAs was marked by yellow trajectories. (G, H) Zoom-up of the highlighted regions in the merge images of (C) and (D). Scale bars: 2 µm.

## SUPPLEMENTARY VIDEOS

**Video S1.**
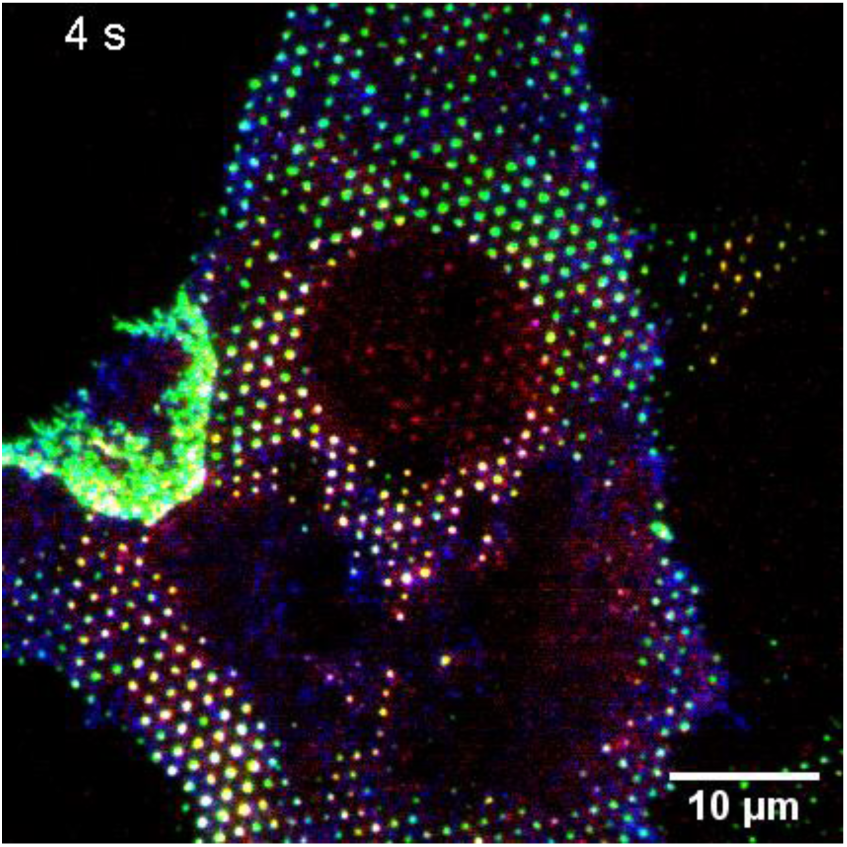
TIRF microscopy of fluorescence recovery after photobleaching of tdmCherry- Axin1(red) on mEGFP-Lrp6 (green)/SNAP-Fzd8(blue) nanodot array. Scale bar: 10 µm.

**Video S2.**
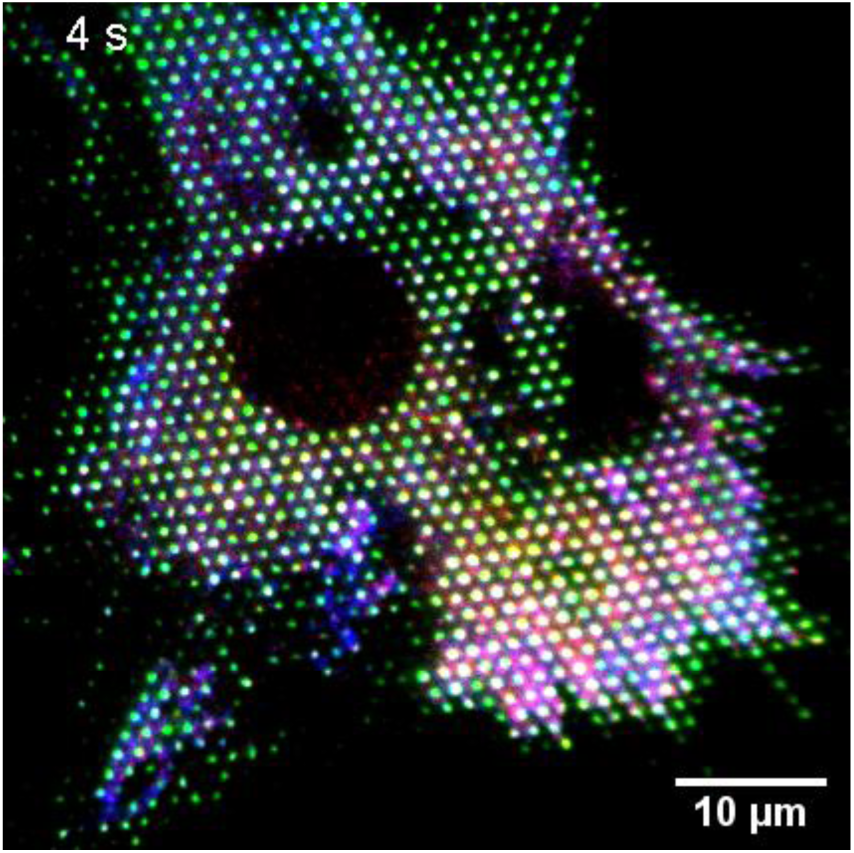
Time-lapse TIRF microscopy of FRAP for quantifying the dynamics of Dvl2-tdmCherry (red) on mEGFP-Lrp6 (green)/SNAP-Fzd8(blue) nanodot array. Scale bar: 10 µm.

**Video S3.**
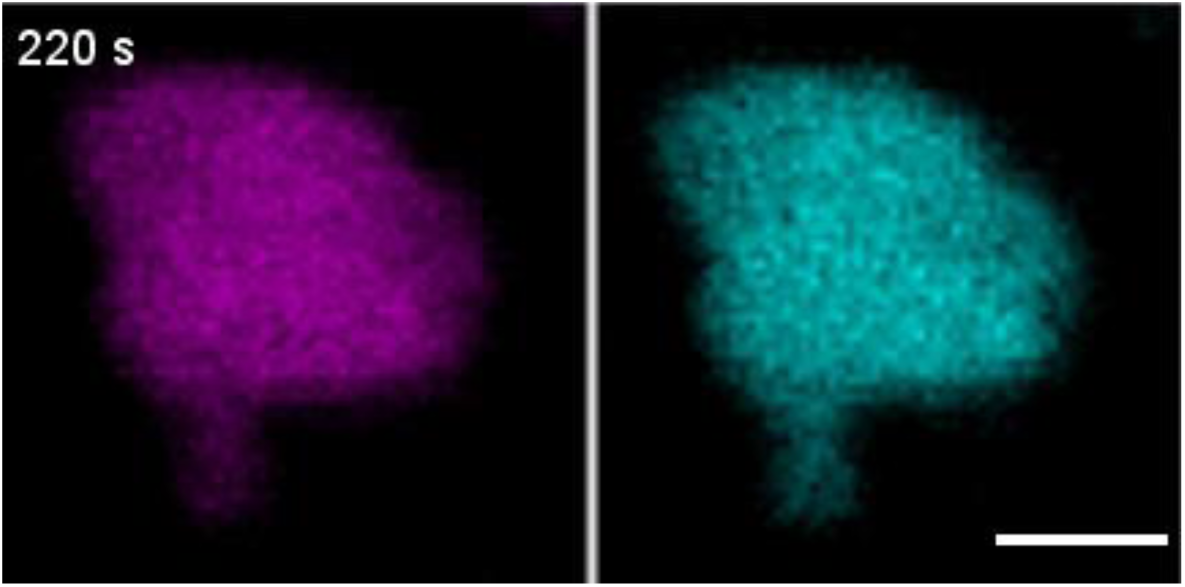
FRAP experiment of a ^SiR^HaloTag-Axin1 (magenta)/Dvl2-tdmCherry (cyan) co-condensate in the cytosol of a HeLa cell. Scale bar: 2 µm.

**Video S4.**
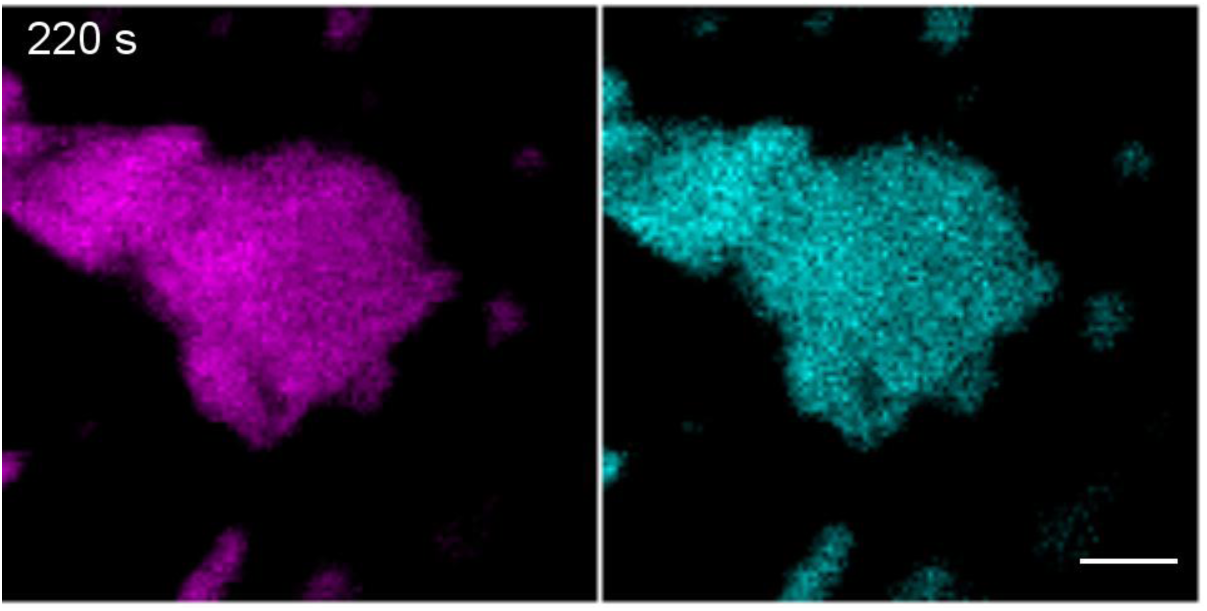
Time-lapse confocal microscopy imaging of sub-droplet FRAP in ^SiR^HaloTag-Axin1 (magenta)/Dvl2-tdmCherry (cyan) co-condensates in a HeLa cell. Scale bar: 2 µm.

**Video S5.**
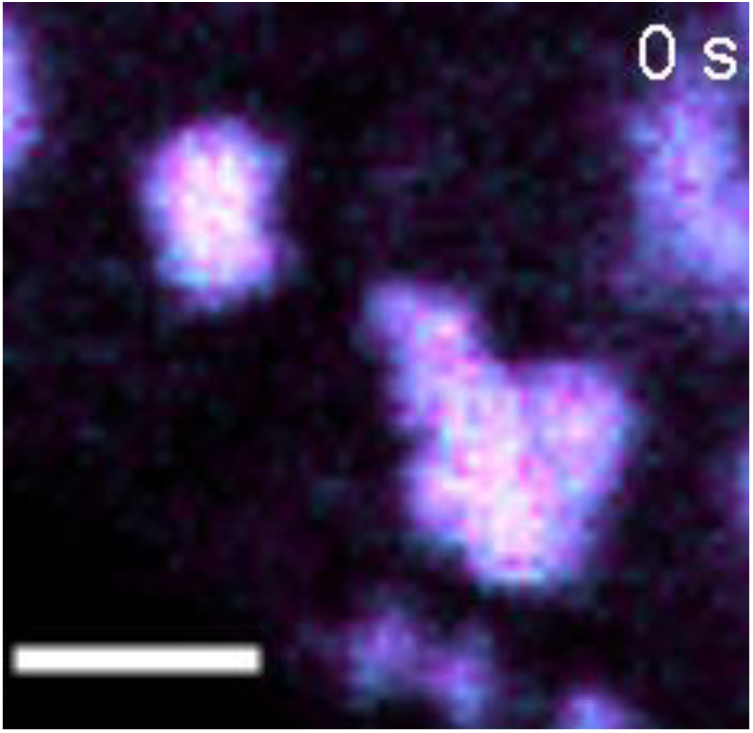
Time-lapse confocal microscopy imaging of the dividing and coalescing of ^SiR^HaloTag- Axin1 (magenta)/Dvl2-tdmCherry (cyan) co-condensates in a HeLa cell. Scale bar: 2 µm.

**Video S6.**
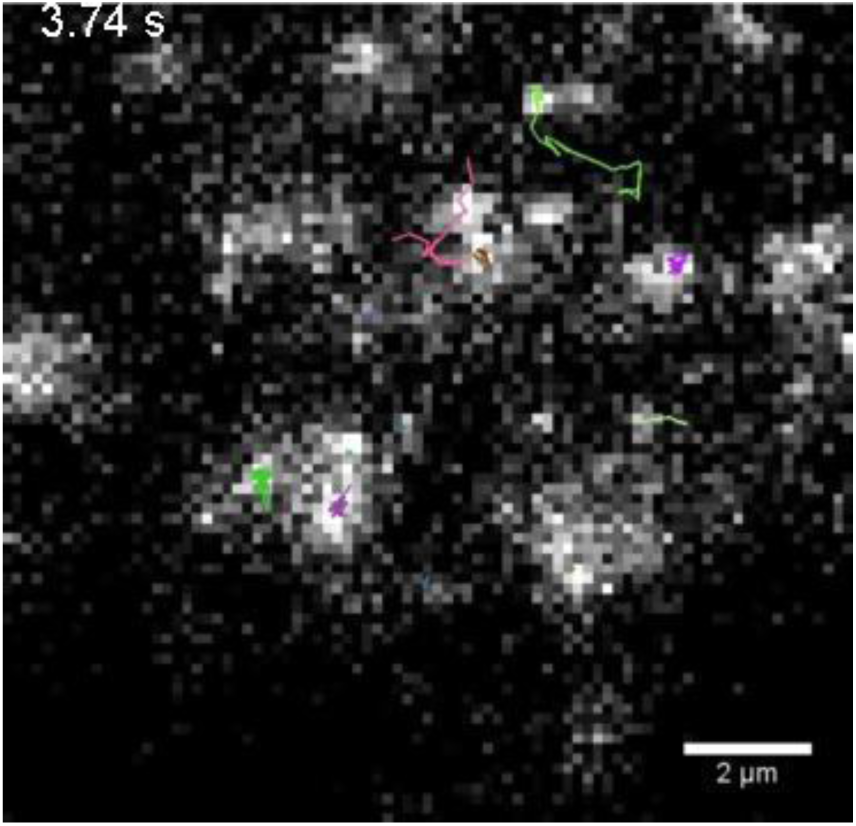
Single molecule tracking of individual Dvl2-HaloTag molecules in Dvl2-^SiR^HaloTag/Dvl2-tdmCherry droplet in a HeLa cell. Scale bar: 2 µm.

**Video S7.**
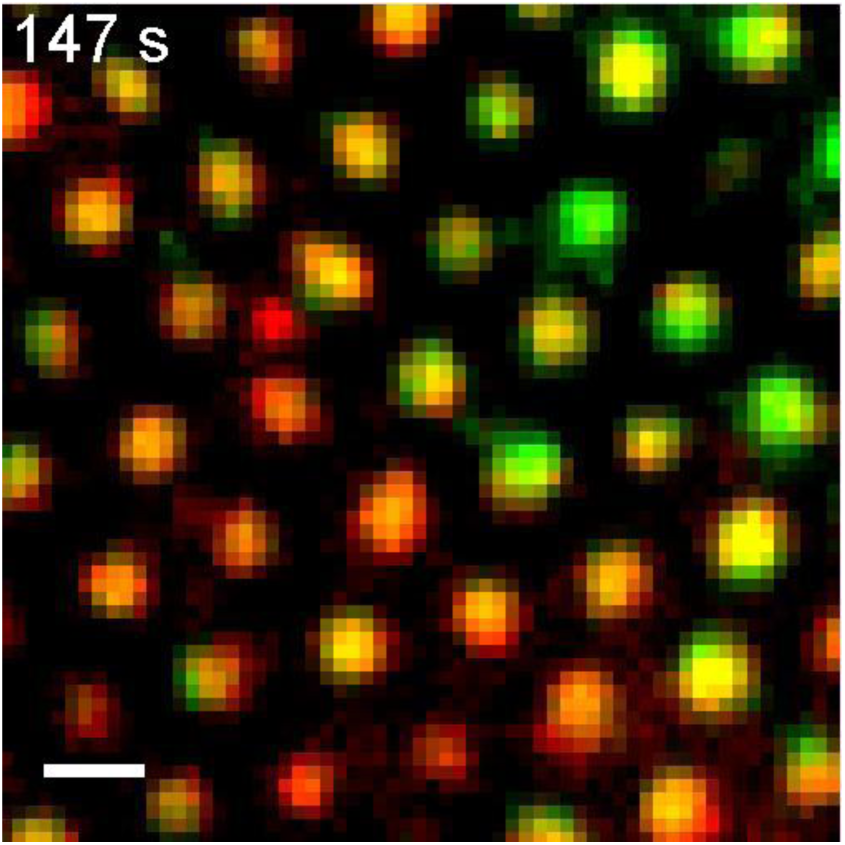
Time-lapse TIRF imaging of tdmCherry-Axin1 nanodroplet arrays (red) generated by direct capturing of Dvl2-mEGFP (green) into bNDAs of an Axin1/2 DKO HeLa cell. Scale bar: 1 µm.

## SUPPLEMENTARY TABLE

**Table S1.** Summary of association rate constants *k_on_* determined by TIRFS-RIF detection.

| Protein | Surface | $k_{on} (M^{-1}s^{-1})^*$ |
| --- | --- | --- |
| BSA | DMDCS | $(1.05 \pm 0.18) \times 10^4$ |
| | OTS | $(9.75 \pm 0.01) \times 10^3$ |
| | DPDCS | $(1.44 \pm 0.03) \times 10^4$ |
| | NMTS | $(2.73 \pm 0.28) \times 10^5$ |
| HaloTag | <sup>HTL</sup>BSA | $(3.26 \pm 0.01) \times 10^3$ |
| mEGFP | tdClamp | $(8.14 \pm 0.08) \times 10^4$ |
| ALFAnb | <sup>ALFA</sup>BSA | $(2.02 \pm 0.05) \times 10^4$ |

\* Data presented as mean ± S.E. of curve fittings.

**Table S2.**
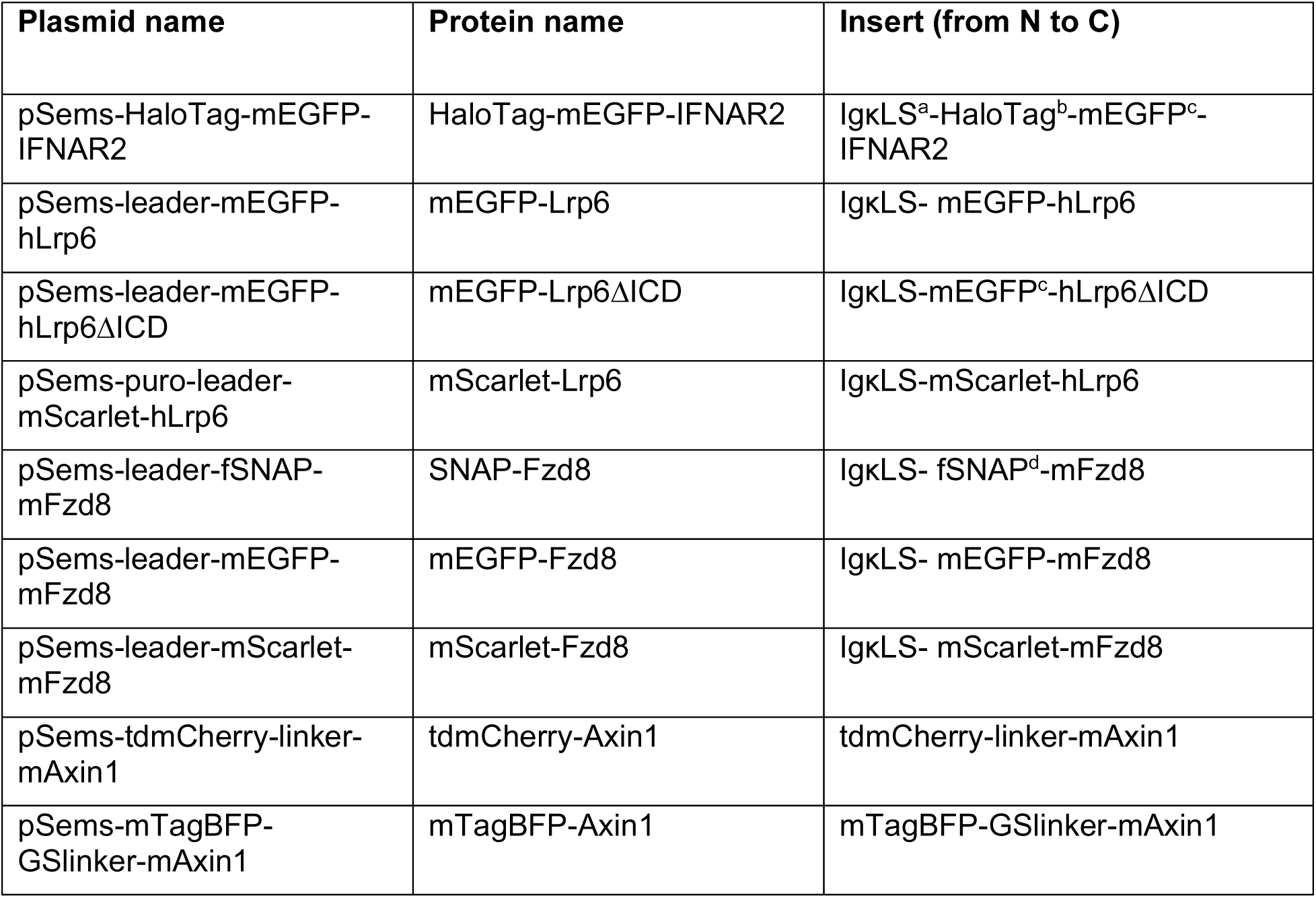
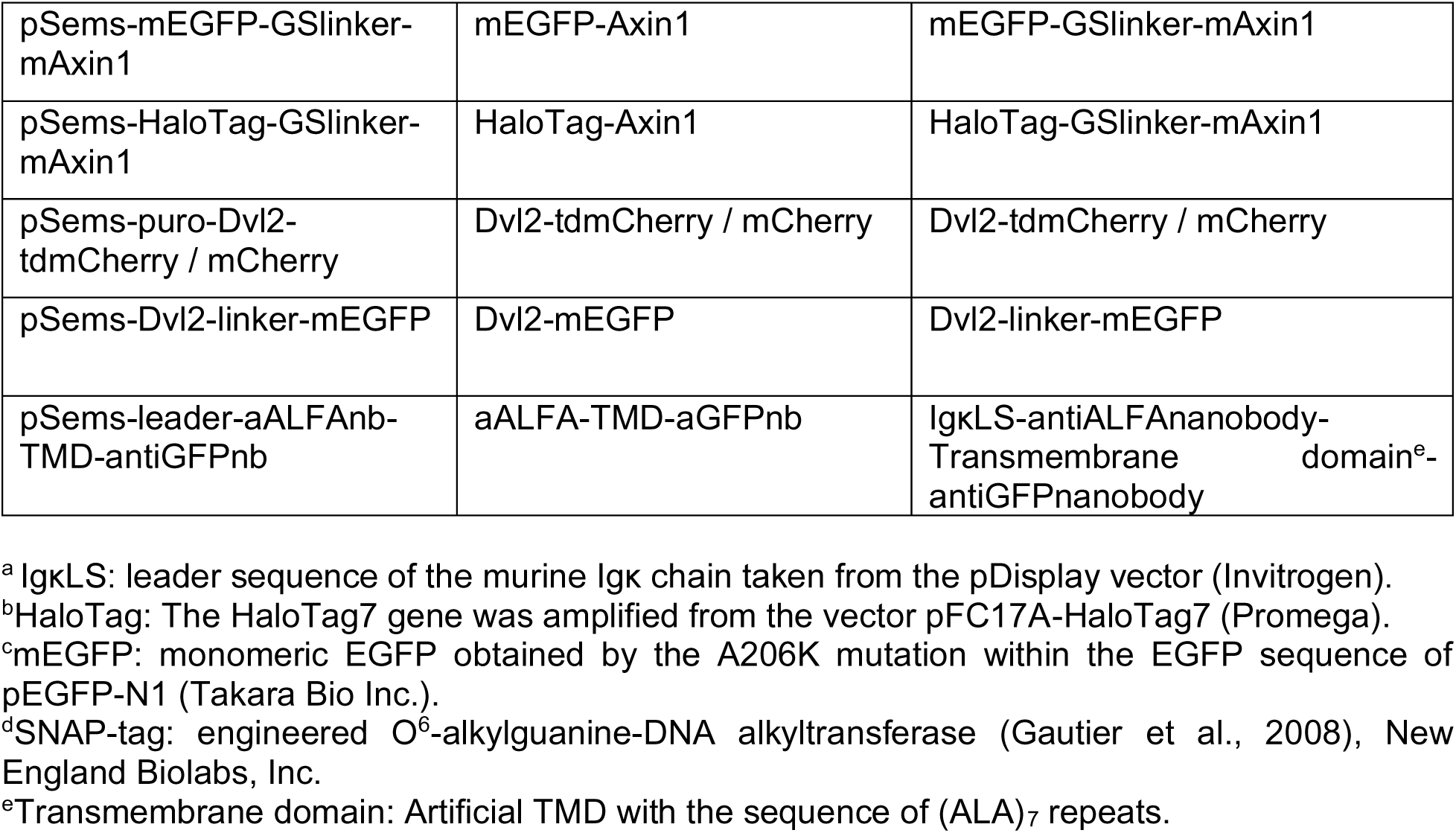
Description of the plasmids used for live cell bNDA experiments.

| Plasmid name | Protein name | Insert (from N to C) |
| --- | --- | --- |
| pSems-HaloTag-mEGFP-IFNAR2 | HaloTag-mEGFP-IFNAR2 | IgkLS<sup>a</sup>-HaloTag<sup>b</sup>-mEGFP<sup>c</sup>-IFNAR2 |
| pSems-leader-mEGFP-hLrp6 | mEGFP-Lrp6 | IgkLS- mEGFP-hLrp6 |
| pSems-leader-mEGFP-hLrp6ΔICD | mEGFP-Lrp6ΔICD | IgkLS-mEGFP<sup>c</sup>-hLrp6ΔICD |
| pSems-puro-leader-mScarlet-hLrp6 | mScarlet-Lrp6 | IgkLS-mScarlet-hLrp6 |
| pSems-leader-fSNAP-mFzd8 | SNAP-Fzd8 | IgkLS- fSNAP<sup>d</sup>-mFzd8 |
| pSems-leader-mEGFP-mFzd8 | mEGFP-Fzd8 | IgkLS- mEGFP-mFzd8 |
| pSems-leader-mScarlet-mFzd8 | mScarlet-Fzd8 | IgkLS- mScarlet-mFzd8 |
| pSems-tdmCherry-linker-mAxin1 | tdmCherry-Axin1 | tdmCherry-linker-mAxin1 |
| pSems-mTagBFP-GSlinker-mAxin1 | mTagBFP-Axin1 | mTagBFP-GSlinker-mAxin1 |
| pSems-mEGFP-GSlinker-mAxin1 | mEGFP-Axin1 | mEGFP-GSlinker-mAxin1 |
| pSems-HaloTag-GSlinker-mAxin1 | HaloTag-Axin1 | HaloTag-GSlinker-mAxin1 |
| pSems-puro-Dvl2-tdmCherry / mCherry | Dvl2-tdmCherry / mCherry | Dvl2-tdmCherry / mCherry |
| pSems-Dvl2-linker-mEGFP | Dvl2-mEGFP | Dvl2-linker-mEGFP |
| pSems-leader-aALFAnb-TMD-antiGFPnb | aALFA-TMD-aGFPnb | IgkLS-antiALFAnanobody-Transmembrane domain<sup>e</sup>-antiGFPnanobody |

<sup>a</sup> IgkLS: leader sequence of the murine Igk chain taken from the pDisplay vector (Invitrogen).
<sup>b</sup> HaloTag: The HaloTag7 gene was amplified from the vector pFC17A-HaloTag7 (Promega).
<sup>c</sup> mEGFP: monomeric EGFP obtained by the A206K mutation within the EGFP sequence of pEGFP-N1 (Takara Bio Inc.).
<sup>d</sup> SNAP-tag: engineered O<sup>6</sup>-alkylguanine-DNA alkyltransferase (Gautier et al., 2008), New England Biolabs, Inc.
<sup>e</sup> Transmembrane domain: Artificial TMD with the sequence of (ALA)<sub>7</sub> repeats.

**Table S3:** List of key materials and suppliers.

| Reagent and abbreviation | Source | Catalogue number |
| --- | --- | --- |
| 1-Napthylmethyl)trichlorosilane (NMTS) | Gelest | SIN6596.0 |
| Diphenyldichlorosilane (DPDCS) | Gelest | SID4510.1 |
| Dichlorodimethylsilane (DMDCS) | Merck | 8034520250 |
| Trichlorooctadecylsilane (OTS) | Sigma Aldrich | 104817 |
| Toluene | Fisher Scientific | 14214914 |
| 1-Ethyl-3-(3-dimethylaminopropyl)carbodiimide-hydrochloride (EDC) | Carl Roth | 2156.2 |
| N-Succinimidyl 13-Maleimido-11-oxo-4,7-dioxa-10-azatridecanoate (Mal-PEG<sub>2</sub>-NHS) | Tokyo Chemical Industry | M3079 |
| Albumin Fraction V from bovine serum (BSA) | Carl Roth | 0163.1 |
| HTL-NH<sub>2</sub> | Own synthesis | Liße, *et al.*, 2011 |
| ALFA-peptide (Ac-CPSRLEEELRRRLTE-COOH) | Romer Labs | Custom synthesis |
| 1,4-Dithiothreitol (DTT) | Carl Roth | 6908.2 |
| N-2-Hydroxyethylpiperazine-N'-2-ethane sulphonic acid (HEPES) | Carl Roth | 6763.3 |
| Ethylenediamine tetra-acetic acid (EDTA) | Carl Roth | 8040.2 |
| Dulbecco's Modified Eagle's Medium (MEM) | PAN-Biotech | P04-09500 |
| Dulbecco's phosphate buffered saline (PBS) | PAN-Biotech | P04-36500 |
| Fetal bovine serum (FBS) | PAN-Biotech | P30-3031 |
| NaCl | Carl Roth | 3957.1 |
| Imidazole | Carl Roth | 3899.4 |
| Isopropyl-β-D-thiogalactopyranoside (IPTG) | Thermo Fisher Scientific | R0392 |
| DNase | Sigma Aldrich | DN25 |
| Lysozyme | Sigma Aldrich | L6876 |
| Urea | Carl Roth | 3941.2 |
| Protease inhibitor | Serva | 39106 |
| Chloramphenicol | Biomol | C4350.25 |
| Ampicilin | Biomol | 01503.25 |
| Trypsin | Capricorn Scientific | TRY-1B |
| ViaFect™ | Promega | E4982 |
| Linear polyethylenimine hydrochloride (PEI) | Polysciences | 24765-1 |
| 2-Amino-2-(hydroxymethyl)propane-1,3-diol (TRIS) | Sigma Aldrich | 8.01464 |
| Catalase | Sigma Aldrich | C-40 |
| Glucose | Carl Roth | 3774.1 |
| Glucose oxidase | Sigma Aldrich | 49180 |
| Glutaraldehyde | Electron Microscopy Sciences | 16216 |
| β-Mercaptoethanol | Sigma Aldrich | 63689 |
| 6-bromoindirubin-3-oxime (BIO) | Sigma Aldrich | B1686 |
| Monoclonal β-Catenin Mouse Antibody – Alexa Flour 647 Conjugate | Cell Signaling Technology | 4627S |
| Silicon Rhodamine HaloTag-Ligand (SiR-HTL) | Kai Johnsson's Lab | Lukinavičius G., *et al.*, 2013 |
| ATTO647N-NHS ester | ATTO-TEC GmbH | AD 647N-31 |
| SNAP-Surface®549 | New England Biolabs | S9112S |
| SNAP-Surface®647 | New England Biolabs | Discontinued and replaced by SNAP-Surface® 649, S9159S |

## Notes

### Competing Interest Statement

The authors have declared no competing interest.

### Summary of Updates

Figure 4 revised. Figures 1 and 6 have minor changes. Discussion expanded to compare the nanopatterning methods. Title updated.

